# Reference plasmid pHXB2_D is an HIV-1 molecular clone that exhibits identical LTRs and a single integration site indicative of an HIV provirus

**DOI:** 10.1101/611848

**Authors:** Alejandro R. Gener, Wei Zou, Brian T. Foley, Deborah P. Hyink, Paul E. Klotman

## Abstract

**Objective:** To compare long-read nanopore DNA sequencing (DNA-seq) with short-read sequencing-by-synthesis for sequencing a full-length (e.g., non-deletion, nor reporter) HIV-1 model provirus in plasmid pHXB2_D.

**Design:** We sequenced pHXB2_D and a control plasmid pNL4-3_gag-pol(Δ1443-4553)_EGFP with long- and short-read DNA-seq, evaluating sample variability with resequencing (sequencing and mapping to reference HXB2) and *de novo* viral genome assembly.

**Methods:** We prepared pHXB2_D and pNL4-3_gag-pol(Δ1443-4553)_EGFP for long-read nanopore DNA-seq, varying DNA polymerases Taq (Sigma-Aldrich) and Long Amplicon (LA) Taq (Takara). Nanopore basecallers were compared. After aligning reads to the reference HXB2 to evaluate sample coverage, we looked for variants. We next assembled reads into contigs, followed by finishing and polishing. We hired an external core to sequence-verify pHXB2_D and pNL4-3_gag-pol(Δ1443-4553)_EGFP with single-end 150 base-long Illumina reads, after masking sample identity.

**Results:** We achieved full-coverage (100%) of HXB2 HIV-1 from 5’ to 3’ long terminal repeats (LTRs), with median per-base coverage of over 9000x in one experiment on a single MinION flow cell. The longest HIV-spanning read to-date was generated, at a length of 11,487 bases, which included full-length HIV-1 and plasmid backbone with flanking host sequences supporting a single HXB2 integration event. We discovered 20 single nucleotide variants in pHXB2_D compared to reference, verified by short-read DNA sequencing. There were no variants detected in the HIV-1 segments of pNL4-3_gag-pol(Δ1443-4553)_EGFP.

**Conclusions:** Nanopore sequencing performed as-expected, phasing LTRs, and even covering full-length HIV. The discovery of variants in a reference plasmid demonstrates the need for sequence verification moving forward, in line with calls from funding agencies for reagent verification. These results illustrate the utility of long-read DNA-seq to advance the study of HIV at single integration site resolution.

## Introduction

Much of what we know about human acquired immunodeficiency syndrome (AIDS) came after isolating the causative agent – the human immunodeficiency virus type 1 (HIV-1) – and describing the viral genome information content. The HIV-1 isolate HXB2 (also known as HTLV-III and HIV-1_LAI_ or LAV/BRU [1], [2]) was the first full-length replication-competent HIV genome sequenced [3]. Derivative clones commonly called “HXB2” are still used for *in vitro* infection assays, including RNA (almost always cDNA [4]) sequencing (**Figure 1A** and **Supplemental Table 1**). Despite the availability of the HXB2 HIV-1 reference sequence [3], no sequence is available for any complete and readily available HXB2 clone.

**Figure 1:**
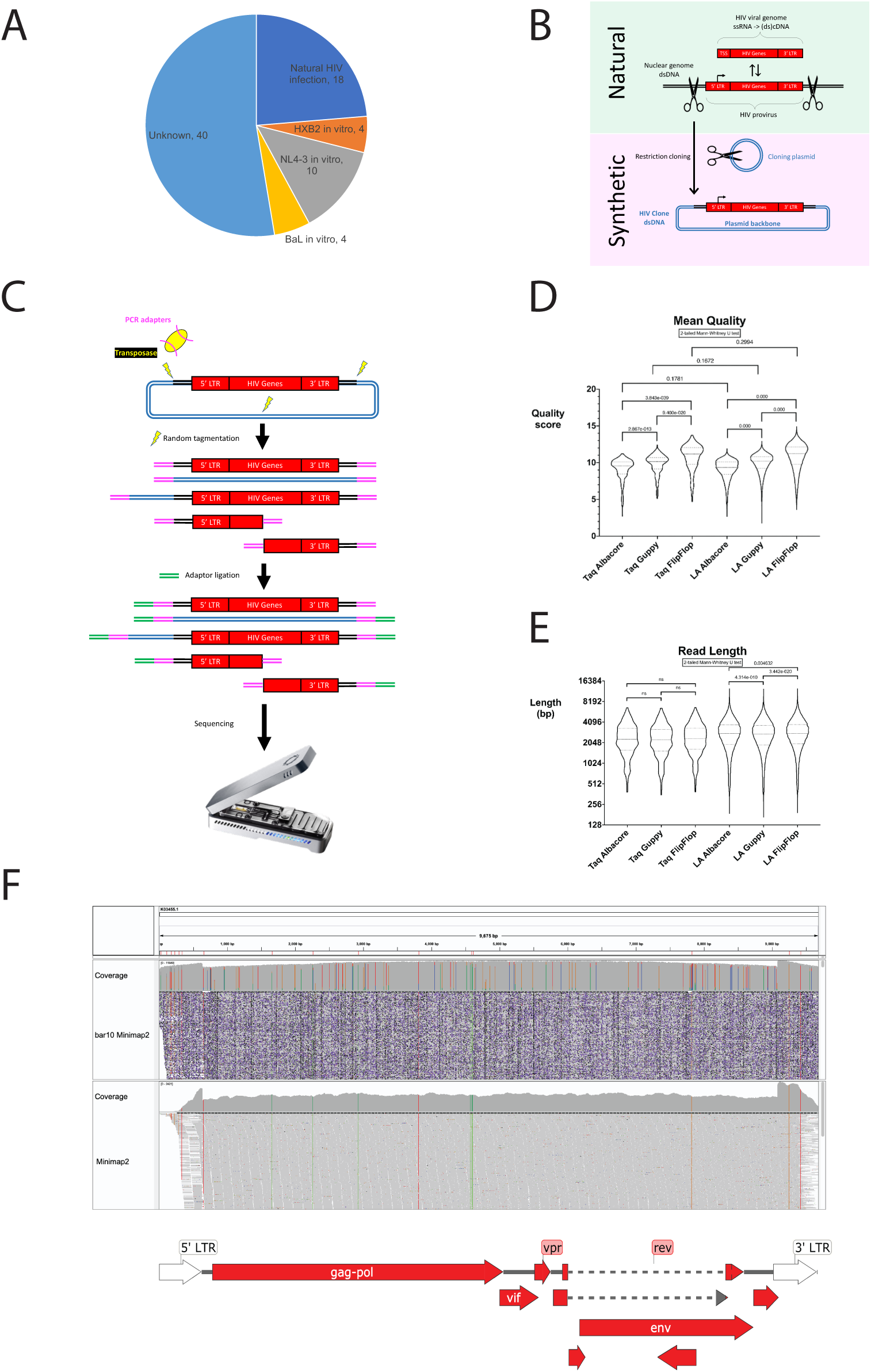
HIV information in pHXB2_D is recovered by long-read sequencing and mapping. Figure 1A: HXB2 is still a commonly used resource. It is the reference HIV-1 genome, derived from one of the earliest clinical isolates. While older HIV samples are occasionally rediscovered, they are not made routinely available to researchers. All public HIV-1 RNA-seq datasets were obtained from the NCBI SRA with the following search phrase: “HIV-1” AND “RNA-seq”. Metadata from these 2527 runs (number current as of 7/21/2020) were used to make a pie chart summary. Figure 1B: HIV information comes from three main sources: proviruses (HIV sandwiched between two assumedly identical full-length long terminal repeats (LTRs)), unspliced HIV mRNAs (also known as viral genomes) starting from the transcription start site and ending in the 3’ LTR [4], and engineered proviruses recovered in their entirety or stitched together from multiple isolates like NL4-3 [18]. Figure 1C: ONT library prep pipeline. Tagmentation cleaves double-stranded DNA, ligating barcoded PCR adapters (magenta). PCR-adapted DNA may be amplified. After amplification and cleanup, ONT sequencing adapters (green) are ligated. Barcoded samples may be pooled and sequenced. Figure 1D: Newer basecallers increase read mean quality. Median (big dash) and quartiles (little dash). Effect of enzyme version was not statistically significant. Figure 1E: Read stats with different callers/aligners. Median (big dash) and quartiles (little dash). Read lengths increase with higher fidelity Taq. Figure 1F: Sequencing coverage with long-vs. short-read single-end 150 bp (trimmed to 142 bp) DNA sequencing. Long-read sequencing covers ambiguously mappable areas missed by short-read in HXB2 reference Genbank:K03455.1 (**Supplemental Figures 3D,3E**), but at the expense of accuracy near homopolymers longer than about 4 nucleobases (**Supplemental Figure 5**). Short-read mapping fails at repetitive elements longer than their read lengths (**Supplemental Figures 3D,3E**). Long read Minimap2 settings: map-ont -k15. Short read Minimap2 settings: Short reads without splicing (-k21 -w11 --sr -F800 -A2 -B8 -O12,32 -E2,1 -r50 -p.5 -N20 - f1000,5000 -n2 -m20 -s40 -g200 -2K50m --heap-sort=yes --secondary=no) (sr).

HIV clones were originally made by choosing non-cutter restriction enzymes to digest intact proviral sequences upstream and downstream of unknown integration sites from infected host cells while sparing HIV-1 sequence, followed by ligation into an *E. coli* cloning vector (plasmid) (**Figure 1B**), allowing for low-error (but not error-free) propagation [5]. These clones became available before tractable sequencing methods permitted routine sequence verification. As such, it was uncommon to sequence them. While funding agencies now require investigators to include in their proposals plans to validate their key reagents, these funders tend to leave the process up to investigators and may not always follow up on whether a given reagent is ever actually validated (or revalidated between changes of hand). Investigators do not regularly validate their clones, in part because there is no universally accepted standard. Instead, a common practice is to assume a given clone, often kindly gifted from a colleague, is as reported. As such, we often do not truly know what we have been working with for 35+ years.

Making sense of the information from HIV sequencing experiments is complicated by many factors, including the cycling that all orterviruses [6] undergo between two major states (as infectious virion RNA and integrated proviral DNA **Figure 1B**), repetitive viral sequences like long terminal repeats (LTRs), non-integrated forms [7], rarity of integration events *in vivo* (reviewed in [8]), and alternative splicing of viral mRNAs [9]. Short-read DNA sequencing (<150 base pairs (bp) in most reported experiments, but up to 500 bp for either Illumina sequencing-by-synthesis or <1,000 bp for chain termination sequencing) provides some information, but analyses require high coverage and/or extensive effort (non-exhaustive examples [10], [11]). These factors limit the ability to assign variants to specific loci within each provirus, as well as at the proviral integration site(s) (reviewed in [12]). Despite progress (HIV DNA) [13], (HIV RNA) [14], [15], [16], researchers have yet to observe the genome of HIV-1 as complete provirus (integrated DNA) in a single read, hindering locus-specific studies. To this end, current long-read DNA sequencing clearly surpasses the limitations of read length of leading next-generation/short-read sequencing platforms. Here we used the MinION sequencer to sequence HIV-1 plasmid pHXB2_D in a pilot study focusing on coverage acquisition (as opposed to full-length sequencing), with the goal of evaluating the technology for future applications.

## Methods

This work did not include human or animal subjects. Nanopore libraries for this work were prepared in their entirety by ARG in a Biosafety Level 2 laboratory on main campus at Baylor College of Medicine (BCM). Nanopore sequencing was completed between April and May of 2018 as two of several control experiments included in the Student Genomics pilot run (**Supplemental Information**). Short-read sequencing was completed in April 2019.

### HIV-1 plasmids

A plasmid, “pHXB2_D” (alternate names pHXB2, pHXB-2D), believed to contain the HIV-1 reference strain HXB2 [17] was acquired from the NIH AIDS Reagent and Reference Program (ARP) via BioServe. pHXB2_D was believed to be a molecular clone (likely a restriction product of HXB2 proviral DNA inserted into an unknown cloning plasmid backbone) from one of the earliest clinical “HXB2” HIV-1 isolates. At the time of this work, it was unknown whether this plasmid was ever sequence-verified before or after the reference sequence for HXB2 was deposited.

The provenance of pNL4-3_gag-pol(Δ1443-4553)_EGFP, a reporter construct of pNL4-3 with a gag-pol deletion between base 1443 and 4553 is known. HIV-1 NL4-3 (pNL4-3) was a fusion of NY5 and LAV/HXB2 plasmids [18] that to our knowledge are not readily available. pEVd1443 [19] was a deletion construct made from pNL4-3 used to make several HIV-1 transgenic animals, including the FVB/N-Tg(HIV)26Aln/PkltJ (The Jackson Laboratory stock No: 022354) “Tg26” mouse. The deletion in pEVd1443 was made by SphI cutting between d1443 and 1444 with binding site 1443-1448, and cutting at a BalI site at 4551-4556 with blunt cutting between 4553 and 4554. The EGFP cassette includes additional sequence upstream and downstream of EGFP coding sequence. SphI and BalI may still be used to excise EGFP cassette. A reporter construct was designed mimicking the pEVd1443 deletion: pNL4-3: ΔG/P-EGFP [20]. Dr. Wei Zou rederived pNL4-3: ΔG/P-EGFP at BCM [21]. Both constructs (plasmid and mouse) retained parts of gag and pol, with limited effects on protein-coding capacity, such as expression of p17 [22]. Based on Addgene naming conventions, we suggest pNL4-3_gag-pol(Δ1443-4553)_EGFP to replace the previous name pNL4-3: ΔG/P-EGFP for clarity.

### HIV-1 reference sequences

The reference sequence of HXB2 is from the National Center for Biotechnology Information (NCBI), Genbank accession number K03455.1. It runs from the beginning of the 5’ LTR to the end of the 3’ LTR, and is 9,719 bp. This is similar to another HIV-1 reference that NCBI uses, AF033819.3. This is a 9,181 base HXB2-like sequence that starts at the 96 bp repeat in the 5’LTR, continues with the 5’UTR (U5), extends past the 3’UTR (U3) to the end of the 96 bp repeat in 3’LTR, with one SNV at the *vpu* start codon aTg to aCg at position AF033819.3:560 or K03455.1:6063. The reference sequence of NL4-3 is as a plasmid with accession number AF324493.1. It runs from the beginning of the 5’ LTR to the end of the 3’ LTR, spanning 9,709 bp, and includes plasmid backbone with total length 14,825 bp.

### Long-read DNA sequencing

A plasmid containing HXB2 was sequence-verified with long-read nanopore sequencing on a MinION Mk1B (Oxford Nanopore Technologies, Oxford, UK). Unless otherwise noted, reagents (and software) were purchased (or acquired) from Oxford Nanopore. Briefly, stock plasmid was diluted to 5 ng final amount in ultrapure water (as two samples) and processed with Rapid PCR Barcoding kit SQK-RPB004 along with 10 other barcoded samples (not discussed further in this manuscript) following ONT protocol RPB_9059_V1_REVA_08MAR2018 (**Figure 1C**), a public description of which is here: https://store.nanoporetech.com/us/sample-prep/rapid-pcr-barcoding-kit.html. Two DNA polymerases were evaluated (barcode 10 used high-fidelity LA (for “long amplicon”) Taq (Takara); barcode 11 Taq (Sigma-Aldrich). Libraries were loaded onto a MinION flow cell version R9.4.1 and a 48-hour sequencing run was completed with MinKNOW (version 1.10.11). Residual reads from subsequent runs were pooled for final analyses. Long read data for pNL4-3_gag-pol(Δ1443-4553)_EGFP was generated in other barcoded experiments (not shown).

Raw data was basecalled (converted from FAST5 to FASTQ format) with Albacore version 2.3.4 (older basecaller), Guppy version 2.3.1 (current official at time of work), and FlipFlop (Guppy development config). Mapping to reference was done with Minimap2 [23] and BWA-MEM [24], implemented in Galaxy (usegalaxy.org) [25]. Alignments (.bam and .bai files) were visualized in the Integrative Genomics Viewer [26] unless otherwise noted. For *de novo* assembly, demultiplexed basecalled reads were fed into Canu version 1.8 [27]. Genome size was estimated to be 16 Kb from agarose gel of undigested, but naturally degraded linearized pHXB2_D (data not shown). SnapGene version 4.3.4 was used to manually annotate contigs from Canu. Blastn (NCBI) was used to identify unknown regions of pHXB2_D. Polishing was performed on ONT-only assemblies with Medaka (https://github.com/nanoporetech/medaka), in Galaxy. Medaka models: r941_min_fast_g303, r941_min_high_g303, r941_min_high_g330. Inference batch size (-b) = 100. The final pHXB2_D assembly and other full-length HIV clones from the ARP were aligned to the most recent human reference genome (hg38) with Minimap2 in Galaxy with the following parameters: Long assembly to reference mapping (-k19 -w19 -A1 - B19 -O39,81 -E3,1 -s200 -z200 --min-occ-floor=100).

### Statistics

Two-tailed Mann-Whitney U tests were used to compare distributions in long-read data. P-values are reported over brackets delineating relevant comparisons. Calculations and graphing were done with GraphPad Prism for macOS version 8.0.2.

### Short-read DNA sequencing

pHXB2_D and control pNL4-3_gag-pol(Δ1443-4553)_EGFP were provided as 35 ul at ∼63 ng/ul to the Center for Computational & Integrative Biology DNA Core at Massachusetts General Hospital, an external DNA sequencing core specializing in high-throughput next generation (short-read) plasmid sequencing and assembly. Neither HXB2/pNL4-3 reference sequences nor pHXB2_D/pNL4-3_gag-pol(Δ1443-4553)_EGFP draft assemblies (from this work) were provided to core staff at the time of sequencing so that testing would remain masked. While the core’s exact library prep is proprietary, multiplexed library prep and 150 single-end Illumina (ILMN) sequencing were most likely performed on a MiSeq with platform-specific reagents (V2 chemistry, per their website) and barcoding. Data was returned as FASTQ. FASTQC [28] was used in Galaxy for in-house data quality control, and read lengths were all 142 bp per this tool. Mapping as above.

### Sequence comparisons

We used MAFFT v7.475 [29], [30] to compare the LTR sequences of pHXB2_D and HXB2, and pNL4-3 and pNL4-3_gag-pol(Δ1443-4553)_EGFP. For cladistics, we used BLAST at HIV-DB (https://www.hiv.lanl.gov/content/sequence/BASIC_BLAST/basic_blast.html) to find other HXB2-like genomes. The top 50 BLAST hits included many sequences pNL43 clones. pNL4-3 is an artificial recombinant of the NY5 clone with LAV and/or the HXB2 clone [18]. The recombination point is marked by an EcoRI restriction site. We then made a multi-sequence alignment with the final pHXB2_D assembly, the top BAST hits, and the HIV-1 M group subtype reference set using GeneCutter (https://www.hiv.lanl.gov/content/sequence/GENE_CUTTER/cutter.html), and built the maximum likelihood tree using IQ-tree (https://www.hiv.lanl.gov/content/sequence/IQTREE/iqtree.html). pNL4-3_gag-pol(Δ1443-4553)_EGFP was not included in the above trees because of absence of divergence from pNL4-3 sequences outside of the EGFP cassette.

## Results

Viewing mapped data in IGV, the long reads (median read length >2000 bp, **Figure 1E**) from both pHXB2_D ONT experiments clearly covered each LTR (**Figure 1F, Supplemental Figures 1**, **3**), while shorter reads collapsed into one of either LTR (**Figure 1F, Supplemental Figures 3D,3E**). This was also seen when long reads were shorter than LTRs (<600 bp). Mappers BWA-MEM and Minimap2 were chosen based on their ability to handle long and short reads. Other mappers were not evaluated. BWA-MEM mapped more ambiguously, piling partially mapped reads between each LTR; Minimap2 mapped with higher fidelity to reference without splitting reads. Coverage as sequencing depth was higher and more even from the higher-fidelity LA Taq library (**Supplemental Figure 1**). pNL4-3 was known to have distinct LTRs because it was a synthetic recombinant. The higher variant density in NL4-3 LTRs enabled mapping and phasing from short-read data only (**Supplemental Figure 2**).

We counted 20 single nucleotide variants (SNVs) in this reference clone of HXB2 (**Table 1, Supplemental Table 3, Supplemental Figure 3E**). These mismatches were seen in all Canu assemblies (**Supplemental Figures 4A,4B**), verified in IGV and/or SnapGene, and were orthogonally verified by short-read sequencing performed by the external core given masked samples (**Supplemental Figure 3E**). These mismatches represent a ∼0.21% divergence from reference HXB2 K03455.1 (20/9719), which was assumed to have perfect identity (0% divergence). Transitions were more common (14/20) (**Table 1**), coinciding with a previous report of increased transitions over transversions in infection models, because transversions are more likely to be deleterious to viral replication (i.e.: to cause protein-coding changes) [31]. Indeed, almost half (9/20) of the observed SNVs occurred in protein-coding regions, even though 92% of HXB2 is coding (791/9719). Of those 9 SNVs in protein-coding regions, 4 caused non-synonymous mutations. One of those occurs in a region overlapping both gag and pol regions, however only pol exhibited a non-synonymous change from valine to isoleucine in p6, at position 2259 relative to HXB2. Other non-synonymous variants occurred at 4609 (in p31 integrase, arginine to lysine), 7823 (in ASP antisense protein, glycine to arginine), and 9253 (in nef, isoleucine to valine). 11/20 SNVs were in LTRs (see **Supplemental Figure 3** for counting based on mapping); 8/20 of these would have been missed with mapping-only variant calling or consensus. The longest HIV-mapping read (**Figure 2**) phased 16/20 SNVs (failed at sites 2,8,10,12, **Table 1**). pNL4-3_gag-pol(Δ1443-4553)_EGFP did not have HIV-1 or plasmid backbone variants supported by long and short reads outside of the EGFP cassette.

**Figure 2:**
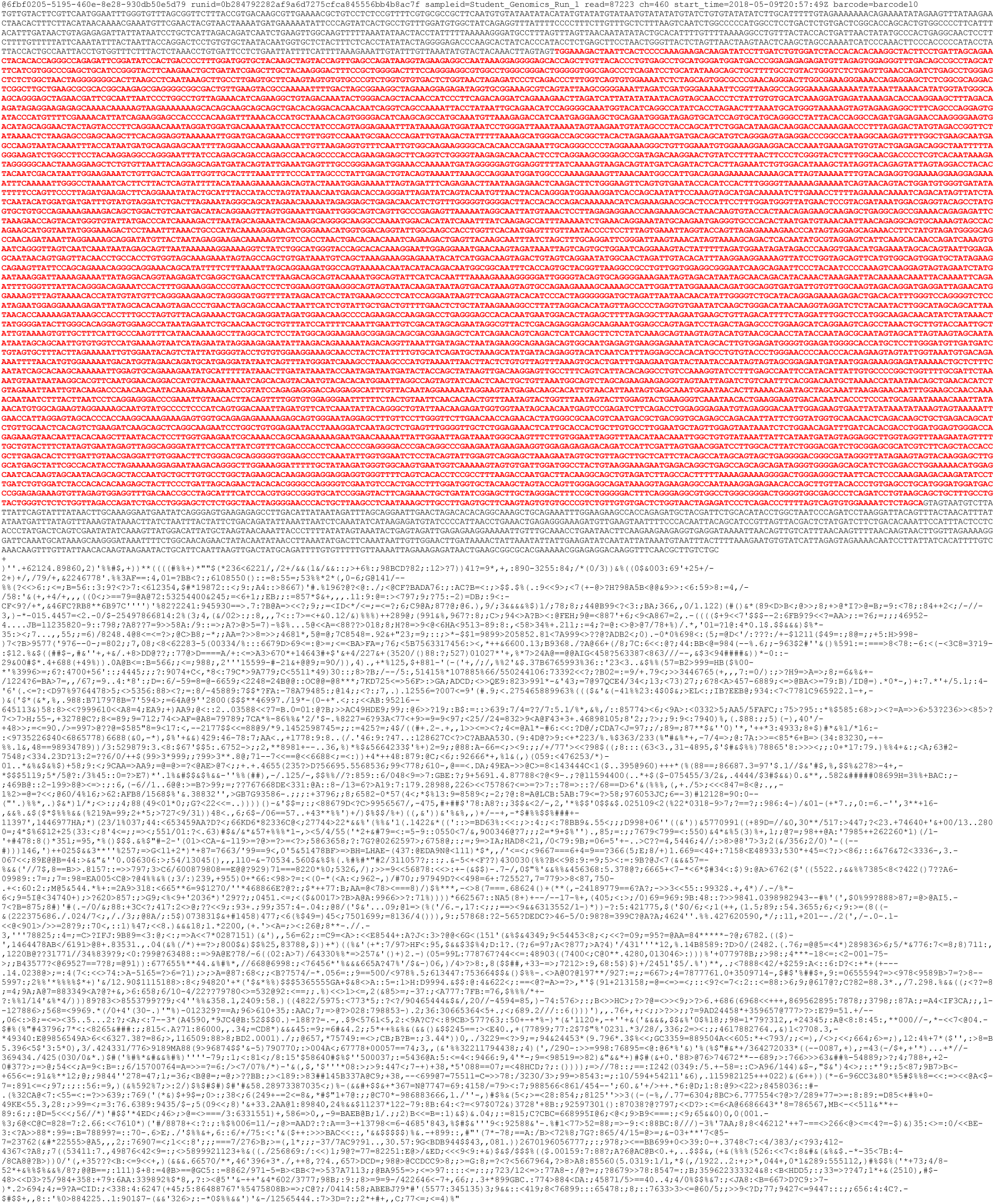
Longest read containing complete full-length HIV-1 reference HXB2. The 5^th^ longest read in the barcode 10 set (read ID 6fbf0205-5195-460e-8e28-930db50e5d79) contained full-length HIV-1. Query (full read) blastn against HIV (taxid:11676) returned 92.95% identity to HIV-1, complete genome (Genbank:AF033819.3). Limiting query to HXB2 (red) blastn against Nucleotide collection nr/nt returned 100% coverage and 93.02% identity to HIV-1 HXB2. This read was 11,487 bases long, with mean quality score 11.984396. Basecalled using Guppy 2.3.1 with FlipFlop config.

**Table 1:**
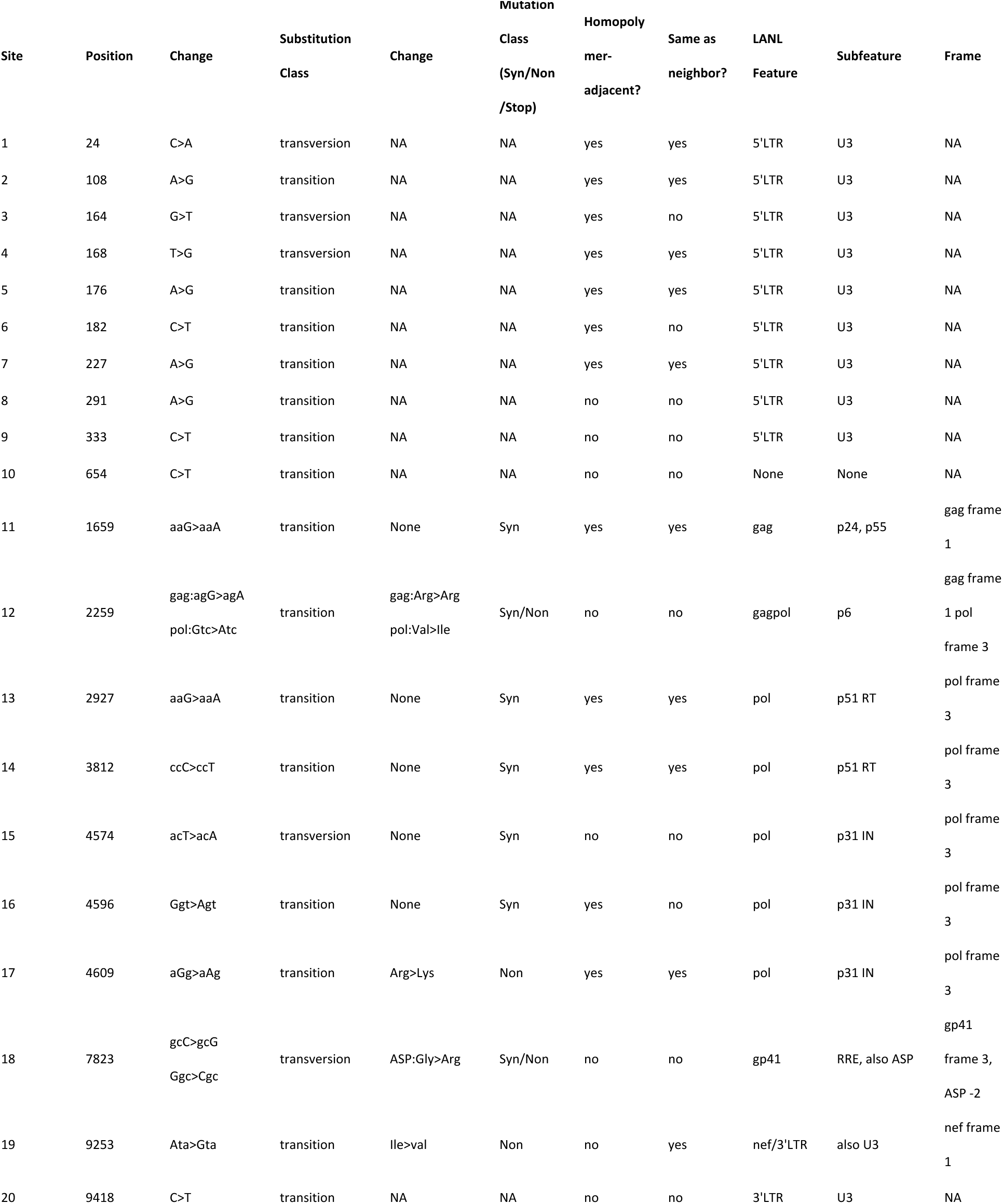
Summary of pHXB2 sample divergence from reference HXB2. Coverage numbers vary by input (albacore, guppy, FlipFlop basecalled FASTQ) and mapping method (Minimap2 vs. BWA-MEM). This information is provided as Supplemental Digital Content. Base-1 (first base is numbered 1, 2^nd^ 2, etc.), relative to HXB2, Genbank:K03455.1. Changed base represented as upper-case. Annotated as codon if in protein-coding region. No deletions or insertions were predicted from manual inspection or supported by short-read sequencing. Abbreviations, ASP: antisense protein, RRE: rev-response element, NA: not applicable. Syn: synonymous mutation. Non: non-synonymous mutation. Stop: stop codon/non-sense mutation. LTR: long terminal repeat. RT: reverse transcriptase. IN: integrase. LANL: Los Alamos National Laboratory HIV Sequence Database. Data from three separate sequencing experiments on the same plasmid sample support these 20 sites. Note site 1-8 variants in 5’LTR have been previously reported (LANL), albethey ambiguously. These may also be incorrectly annotated as variants in nef.

We assembled the previously undefined plasmid pHXB2_D (**Supplemental Figures 4A,4B**). Canu’s final output was a set of contiguous DNA sequences (contigs) as FASTA files. A consequence of assembling plasmid sequences with this tool was partial redundancy at contig ends (**Supplemental Figure 4C)**. Manual end-trimming of contigs was performed in SnapGene based on an estimated length of 16 kilobases. Top blastn hits from barcode 10/LA Taq pHXB2 basecalled with FlipFlop were as follows: for the main backbone (with origin of replication and antibiotic selection cassette for cloning), shuttle vector pTB101-CM DNA, complete sequence (based on pBR322), from 4352-8340; for the upstream element (relative to 5’ LTR), Homo sapiens chromosome 3 clone RP11-83E7 map 3p, complete sequence from 58,052 to 59,165; for the downstream element, cloning vector pNHG-CapNM from 10,204 to 11,666. Other identified elements included Enterobacteria phage SP6 (the SP6 promoter, per SnapGene’s “Detect common features”), complete sequence from 39,683 to 39,966. Identities of query to HXB2 and hits were all approximately 99%. The MGH CCIB DNA Core’s proprietary *de novo* UltraCycler v1.0 assembler (Brian Seed and Huajun Wang, unpublished) was able to assemble both 5’ and 3’ LTRs with short-read data only but may have collapsed SNVs into an artificial single consensus. Long-read mapping and assembly (and polished assemblies) orthogonally validated LTRs, and supported a single HIV-1 HXB2_D haplotype (**Supplemental Figure 4,6**). A final LTR-phased and annotated assembly leveraging short and long reads is provided as pHXB2_D Genbank:MW079479 (embargoed until publication). Importantly, for pHXB2_D, each LTR was identical, which is distinct from the current HXB2 (K03455.1) (**Figure 3A**). Compared to pNL4-3_gag-pol(Δ1443-4553)_EGFP (ACCESSION_TBD) , each LTR was distinct, but identical to pNL4-3’s distinct 5’ and 3’ LTRs (AF324493.1) (**Figures 3B,6**).

**Figure 3A:**
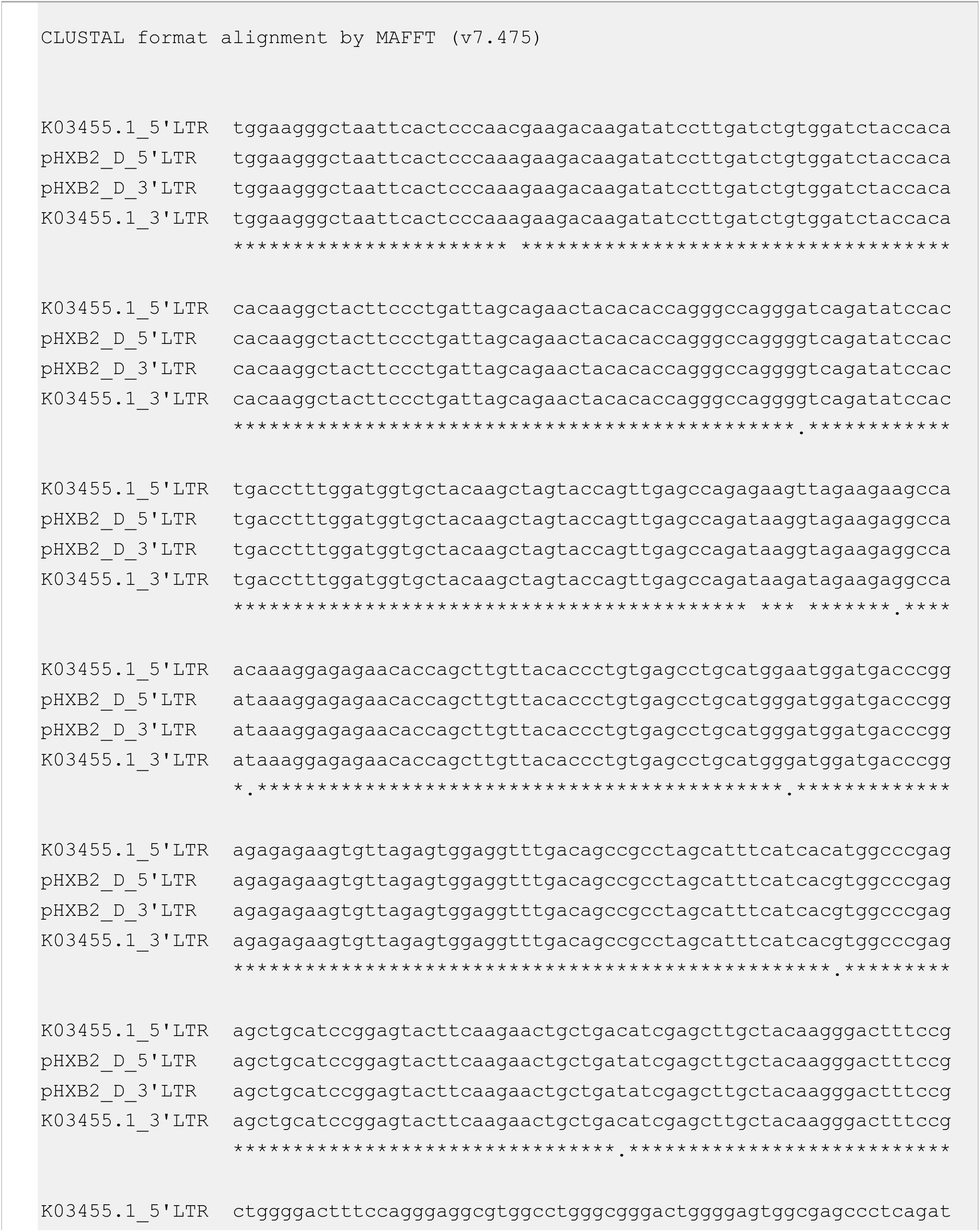
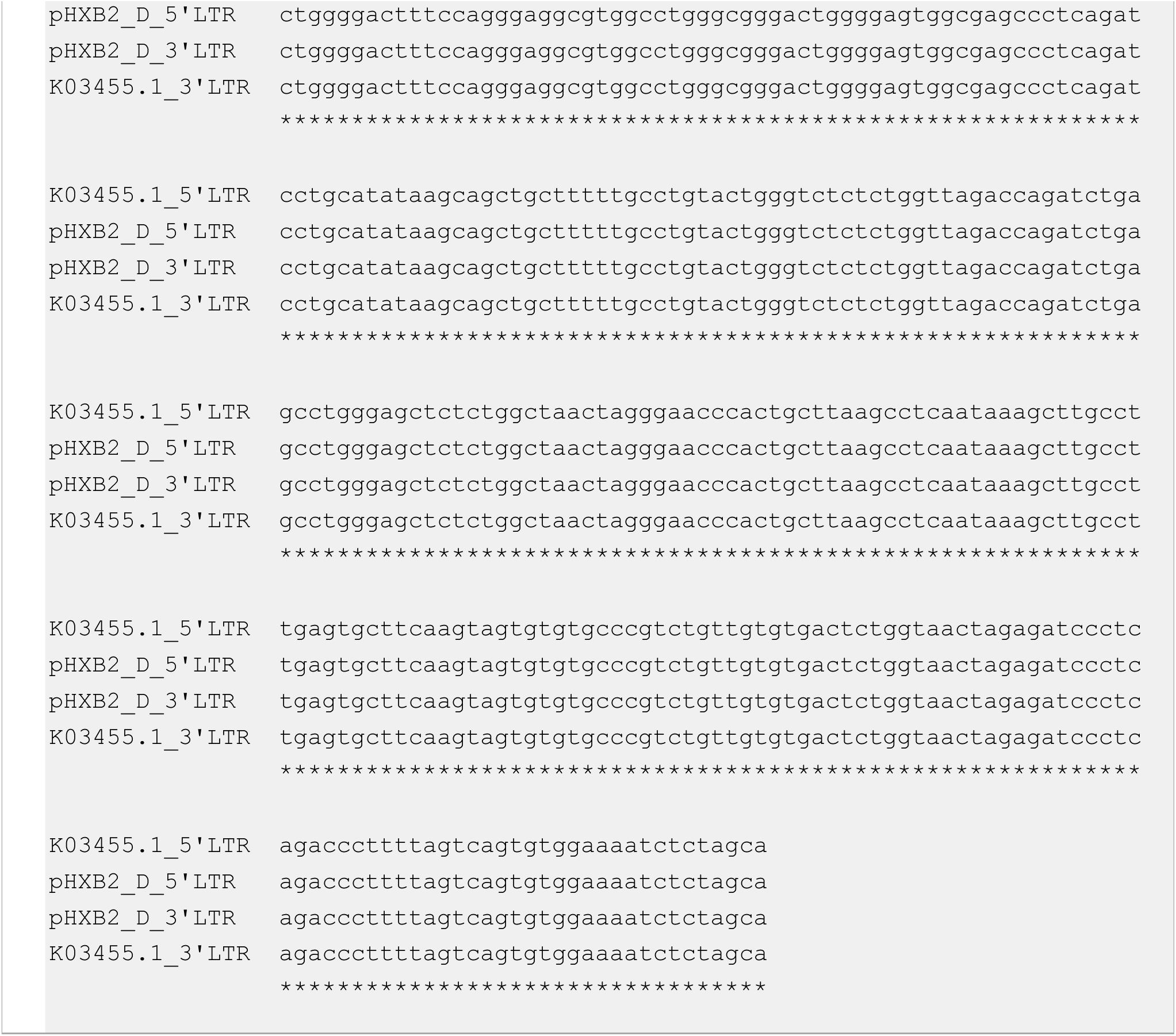
pHXB2_D has identical LTRs, resolving likely errors in HXB2 (K03455.1)

**Figure 3B:**
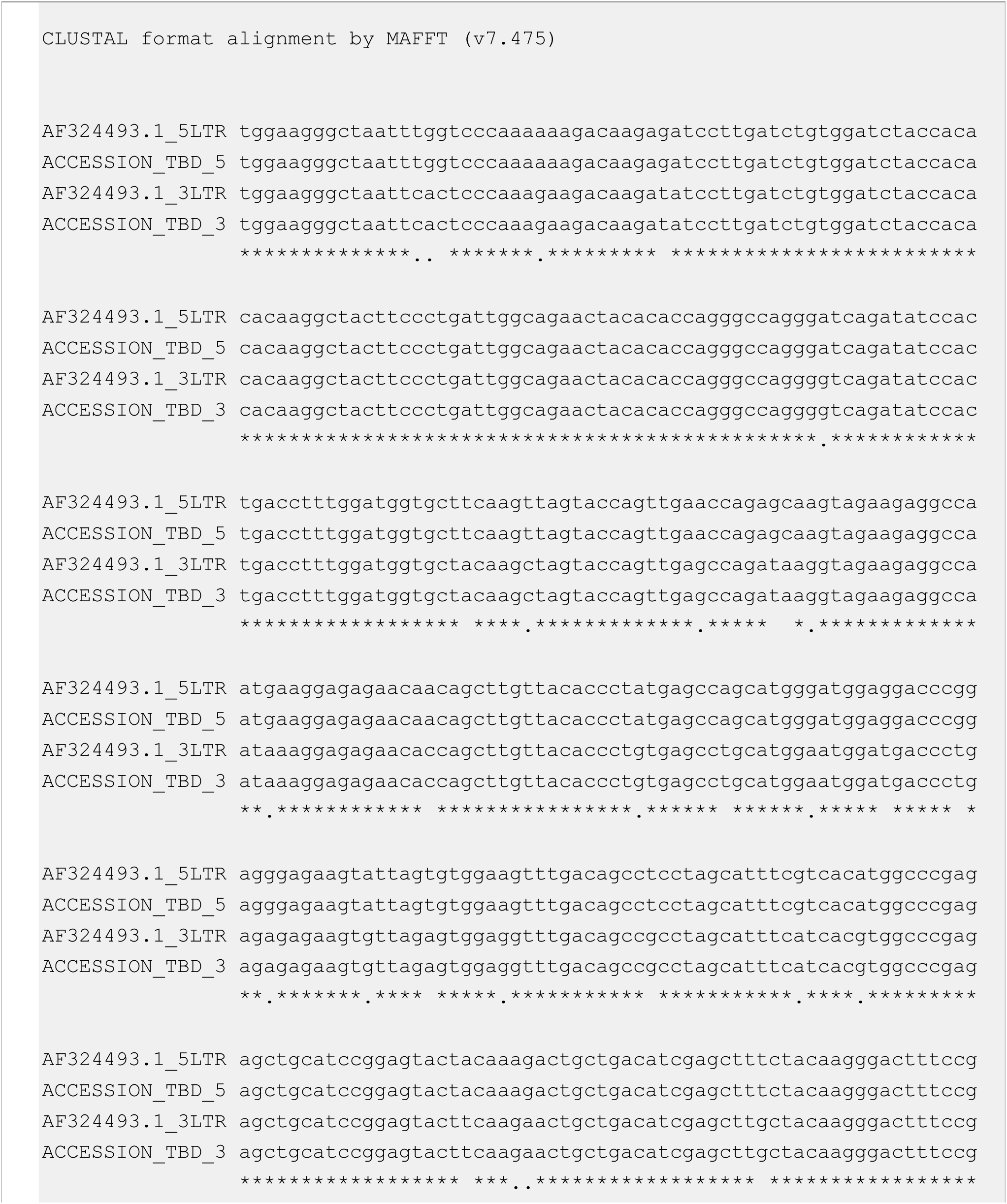
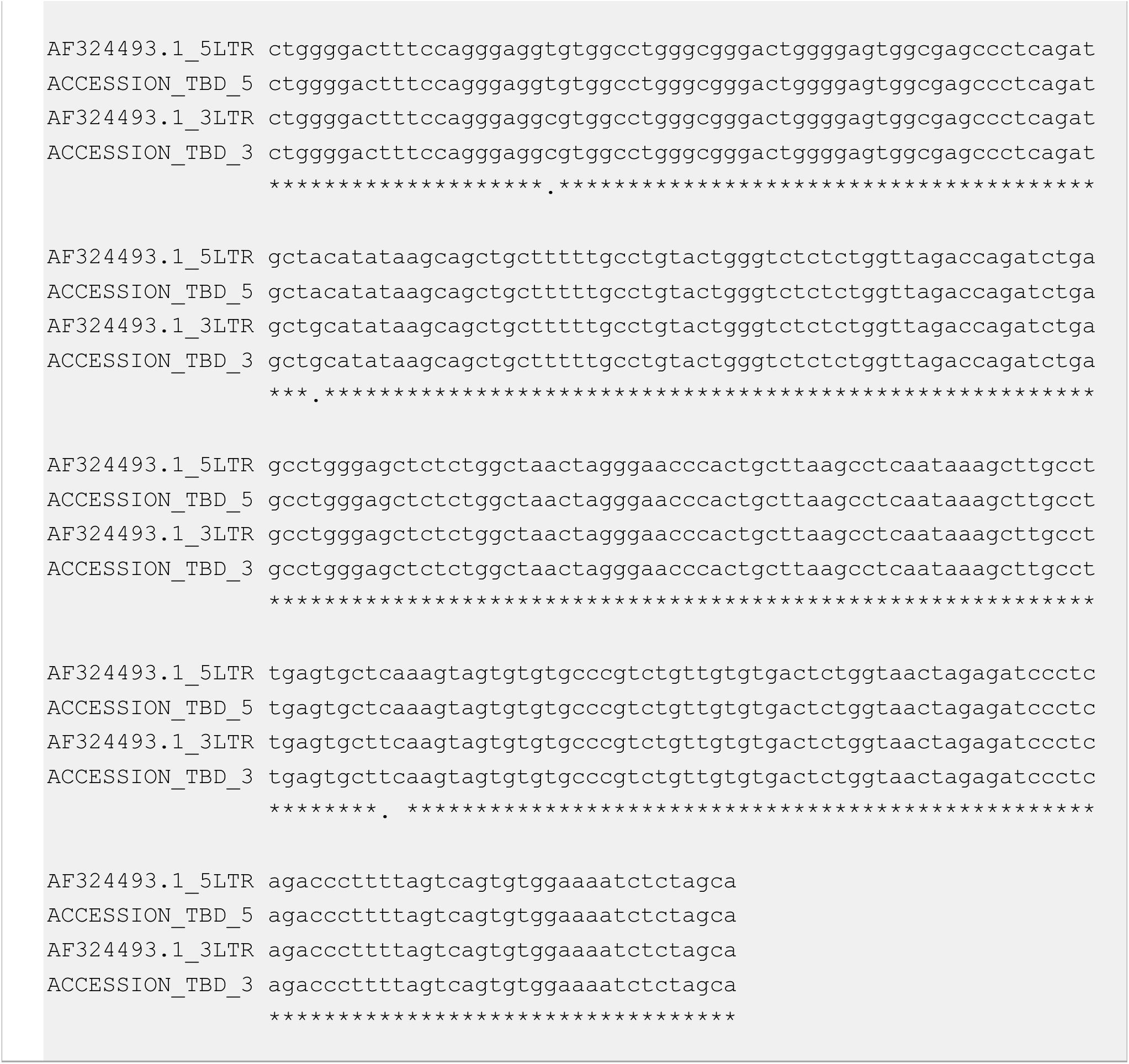
pNL4-3_gag-pol(Δ1443-4553)_EGFP (ACCESSION_TBD) has distinct LTRs, consistent with pNL4-3 (AF324493.1)

To determine whether pHXB2_D was an isolated provirus (as opposed to a cDNA clone), the pHXB2_D assembly was aligned to the current human reference hg38, returning a single complete insertion site on 3p24.3 (**Figure 4A, Supplemental Table 2**). As expected, our pNL4-3_gag-pol(Δ1443-4553)_EGFP had homology arms from two chromosomes (**Figures 4B,6, Supplemental Table 2**). We sought to put our pHXB2_D assembly into context of other HXB2-like references available (**Figure 5**). pHXB2_D (red) clusters closely with HXB2 reference (K03455) and related clone sequences (green). pNL4-3 clones in blue. The LTR-masked HIV-spanning segment of pHXB2_D is most homologous to B.FR.1983.DM461230 and B.FR.1983.CS793683, which are identical except for areas in nef and a GFP insertion (verified by blastn). This finding suggests they were from the same stock. HIV-1 M group subtype reference set (HIV Sequence Database) was added to put HXB2s and pNL4-3 clones into perspective. HXB2 (believed to be a complete isolate) and NL4-3 (synthetic clone based on two early isolates [18]) are examples of HIV type 1 (HIV-1), group M, subgroup B.

**Figure 4A:**
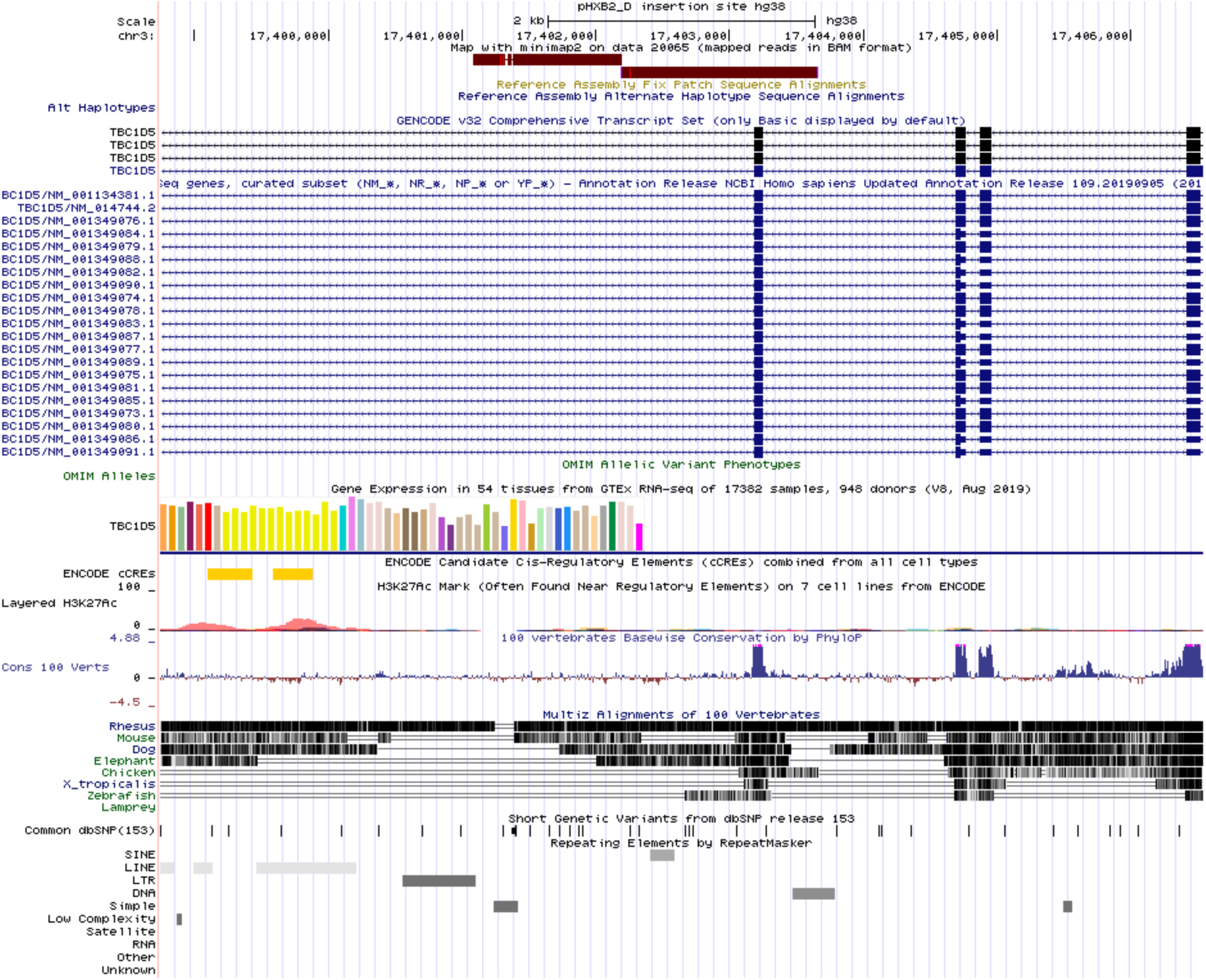
HXB2 integration site

**Figure 4B:**
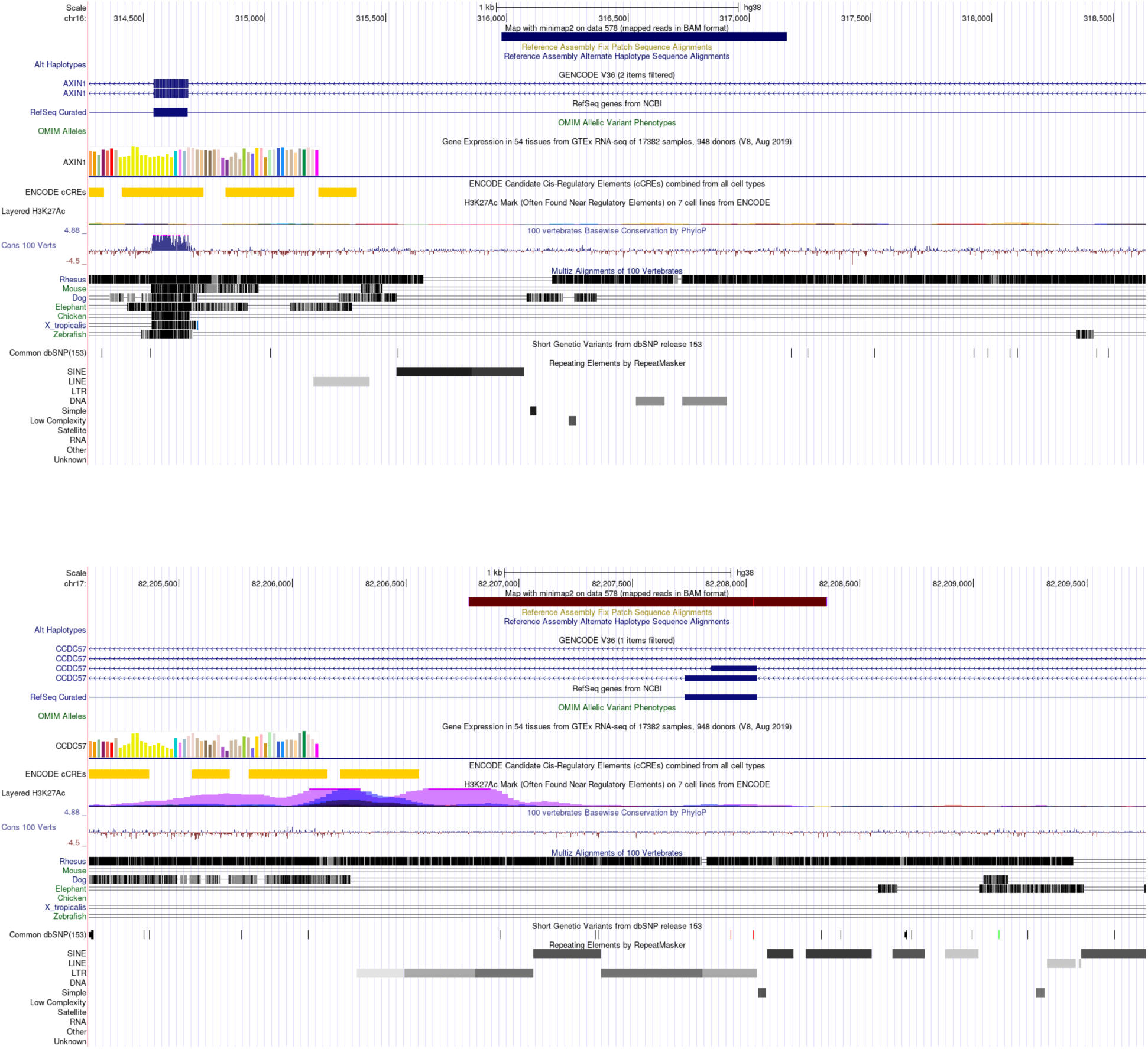
NL4-3 integration sites. Figure 4A: pHXB2_D’s, and therefore HXB2’s, integration site is unambiguously singular (falls outside of annotated repeat), and in the same orientation (minus strand relative to hg38) as target gene TBC1D5. Alignment quality is 60 for both homology arms (**Supplemental Table 2**). Features captured by homology arms in pHXB2_D and other clones verified as proviruses in the present study are consistent with HIV-1 integration behavior [44]. Visualized in UCSC Genome Browser [45]. Figure 4B: pNL4-3_gag-pol(Δ1443-4553)_EGFP’s, and therefore NL4-3’s, integration sites fall on annotated repeats, the longer reads help to locate both sites. Alignment quality is 60 for both homology arms (**Supplemental Table 2**). These integration sites would likely be missed by any method leveraging reads shorter than the homology arms.

**Figure 5:**
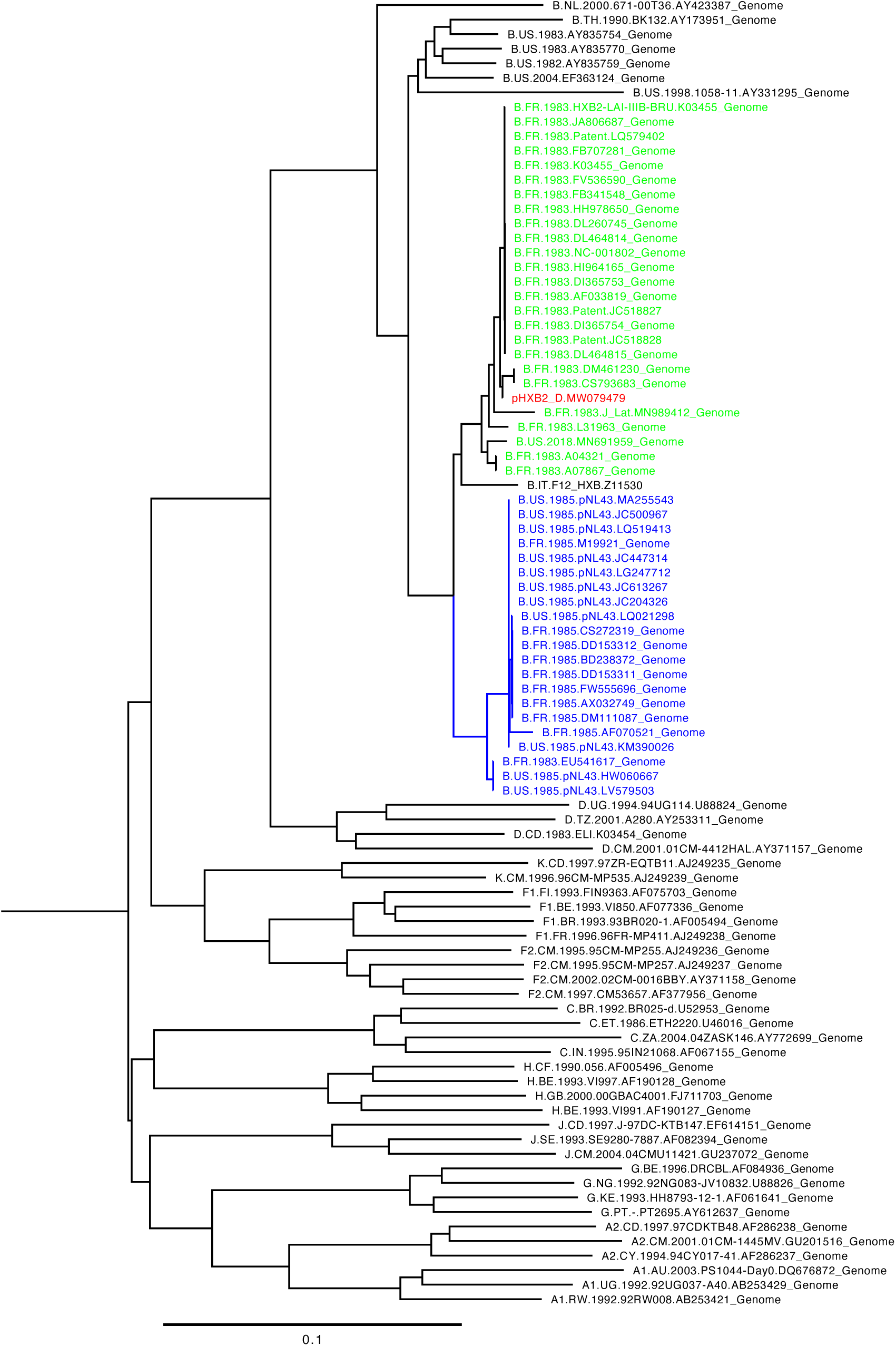
pHXB2_D provenance and top 50 neighbors.

As previously reported [32], per-read variability in ONT data was higher near homopolymers (runs of the same base) (**Supplemental Figure 5A**). For the datasets generated in the present study, homopolymers were counted and classified as continuous (unbroken run of a given nucleobase) vs. discontinuous (broken run of a given nucleobase) (**Supplemental Figures 5B,5D,5F,5H**). A/T (2 hydrogen bonds; 2H) and G/C (3 hydrogen bonds; 3H) were evaluated. Because runs longer than 4 or 5 were rare in these datasets, it was impossible to evaluate longer homopolymers. A simple calculation Abs(Δ)=Abs(#homopolymers_reference_ -#homopolymers_assembly_) helped to evaluate the performance of basecallers, such that better basecallers had smaller Abs(Δ) (**Supplemental Figures 5C,5E,5G,5I,5K**). At the level of consensus (made from sequences mapped to reference HXB2), homopolymers contributed few, if any, obvious errors. A special case of homopolymer, dimer runs, was noted to cause persistent errors regardless of ONT basecaller (**Supplemental Figures 5J,5K**). While dips occurred at certain points near homopolymers, the consensus did not change much at the sequencing depth used in this study for either barcoded pHXB2_D samples (**Supplemental Figures 1,3,4**). Another interpretation is that homopolymers tend to seem truncated with ONT, with more reads in support of shorter homopolymers. Canu assemblies showed basecaller-dependent variability (**Supplemental Table 3**). That said, newer basecallers tended to produce fewer and smaller per-read truncations. Assemblies without polishing did not correct all homopolymer truncations (**Supplemental Figure 4A**). Polishing assemblies tended to correct these toward the final pHXB2_D assembly (**Supplemental Figures 4B,6**). Data from polished ONT-only assemblies and short-read sequencing do not support the truncations (gaps relative to reference) suggested by unpolished ONT-only assemblies, representing a known current limitation of ONT. These are not the same as the 20 SNVs supported by BOTH long- and short-read sequencing performed in this study. The ratio of per-read deletions to per-read insertions (DEL/INS) was much higher for SNVs occurring at homopolymers and near the same base, and this difference was maintained between all basecallers used (**Supplemental Figure 5L**). These changes created more problematic (longer) homopolymers.

## Discussion

This work represents the first instance of complete and unambiguous sequencing of HIV-1 provirus as plasmid and contributed to the identification of single nucleotide variants which may not have been easily determined using other sequencing modalities, illustrating the importance of validating molecular reagents in their entirety, and with complementary approaches. Nanopore sequencing surpassed the read length limitations of traditional sequencing modalities used for HIV such as Sanger sequencing and sequencing-by-synthesis by at least two orders of magnitude. Other long-read DNA sequencing technologies such as PacBio’s zero-mode waveguide DNA sequencing were not evaluated in this work, but in principle would be interchangeable for nanopore sequencing. Paired-end sequencing (as either DNA-seq or RNA-seq) was not evaluated in this work, but has shown promise phasing LTRs in our hands [33]– [35].

### First complete pass over all HIV information in reference plasmid pHXB2_D

HIV provirus is believed to occur naturally as one or a few copies of reverse-transcribed DNA forms integrated into the host nuclear genome. Depending on where integration occurs, local GC or AT content might cause problems for detecting integrants with PCR. HIV also has conserved transitions from areas of higher GC content (∼60%) to content approximating average human GC content (∼40%). To limit PCR sequencing bias and to accommodate for the potential heterogeneity of HIV sequences, we fractionated whole sample directly (as opposed to PCR-barcoding select amplicons) with tagmentation provided in the Rapid PCR-Barcoding kit (ONT). Tagmentation in this workd used transposon-mediated cleavage and ligation of barcode adapters for later PCR amplification. A consequence of this fractionation was a distribution of reads (**Figure 1E**) shorter than longer reads reported elsewhere for ONT experiments [36]. Based on this distribution and the level of coverage, it was expected that HIV might be covered from end to end, but this would have been exceptional. That said, an example is presented here (**Figure 2**). The provirus status of pHXB2_D is supported by recovery of both upstream and downstream homology arms which map to a single human integration site.

### Long reads enable LTR phasing and HIV haplotype definition

We created 6 assemblies for pHXB2_D from ONT-only data (**Supplemental Figure 4**), each with a common set of 20 SNVs (11 in LTRs), and final assemblies (a single HIV-1 HXB2_D haplotype; a single HIV-1 NL4-3_gag-pol(Δ1443-4553)_EGFP haplotype) leveraging long- and short-read data. The external core’s *de novo* assembly pipeline identified the same 20 SNVs, and variants in the LTRs were supported by ONT unambiguously. That the core’s assembler was able to phase LTR variants in these samples may have been because the samples had high amounts of the same upstream and downstream sequences because of coming from one plasmid. The core’s assembler thus may have had additional sequencing information at the edges of HXB2, helping it to map deeper into each LTR. This approach would likely fail in samples with multiple integrations (as in various animal models of HIV disease [37]), which have unknown upstream and downstream sequences, or in samples from natural human infection, which is well known to exhibit multiple pseudo-random integration sites between cells [38], [39], but with mostly single integration events per cell [8]. Inverse PCR (iPCR) is an alternative method [40] with its own issues (e.g., PCR biases, HIV concatemers, host repeats). While current PCR reagents have extended the range of what can be seen with iPCR, current approaches are likewise limited by long DNA extraction methods, sample amount, and remain to be optimized. If coverage is sufficient (≥10 reads in non-homopolymers and non-dimer runs), long-read sequencing can provide linked variant information to individual integration sites. Identical 5’ and ‘3 LTRs (**Figure 3**) in the context of a single integration event (**Figure 4A**) support this integrant being a *bona fide* provirus [41]. Other proviruses also had identical LTR pairs (**Supplemental Table 2**). Technical limitations such as PCR errors before earlier sequencing may explain the variability in the HXB2 reference LTRs. These were sequenced at a time before paired-end 150 or long-read DNA-seq were available to phase LTRs, raising the possibility that these LTRs were incorrectly annotated by depositors assuming identity and copy-and-pasting the sequence of one LTR for both without being able to unambiguously resolve each LTR.

### Mutations in a reference HIV-1 plasmid illustrate the need for reagent verification

Up until 2020, HIV had been the most studied human pathogen, but HIV reagents are not routinely re(verified). The pHXB2_D sequenced was allegedly a reference plasmid, with unknown divergence between the published reference HXB2. Three independent experiments (two long-read with PCR-barcoded libraries made with regular and long-amplicon Taq master mixes, one short-read) yielded at least 20 single nucleotide variants in pHXB2_D which differed from the HXB2 reference sequence (**Table 1, Supplemental Figure 3**), which were also concordant across the three basecallers used (**Supplemental Table 3**) and are therefore not PCR errors. By leveraging long reads with the MinION, we were able to find mutations in highly repetitive LTRs relative to HXB2 Genbank:K03455.1 which are often assumed (but until now never proven) to be identical (**Table 1**, **Figure 1**, **Supplemental Figures 1, 3E**), as well as mutations in protein-coding regions (**Table 1**). We were also able to confirm that the backbone of this plasmid is from pSP62 [17], a pBR322 derivative with the SP6 promoter [42], aiding in the continued use of this important reagent, and illustrating the need of full-length reagent validation moving forward. We suggest that all clinical reagents (e.g., vectors) be sequence-verified at the level of single-molecule sequencing as standard quality control to protect against sample heterogeneity.

### Improvement in ONT basecallers over time

Albacore, Guppy, and FlipFlop basecallers were compared. Each produced reads of similar length distributions (relative to polymerase used), while Guppy and FlipFlop produced improved and best performance relative to quality score distributions (**Figure 1D**). Interestingly, while read length distributions were affected by fidelity of polymerases evaluated in this work, mean quality distributions were not. This is important because of the differences in cost between higher fidelity Taq and classic Taq enzymes. That said, higher fidelity LA Taq produced much higher coverage compared to Taq (**Supplemental Figure 1**). In consideration of library prep, choice of enzyme used should be based on the desired read-length distribution and coverage. Regarding read mapping, the increase in mean quality score between these basecallers improved overall mapping, in part by facilitating demultiplexing, resulting in approximately ∼10% increases number of reads in barcoded libraries before mapping (shift in reads from unclassified to a given barcode). FlipFlop tended to handle homopolymers better than previous basecallers (**Supplemental Figures 5,6**). Homopolymers in HXB2 tended to exhibit apparent deletions near 5’ ends of homopolymers (upstream due to technical artifact from mapping), but because consensus is conserved (example, at least 80% of base in called read set is identical to reference), and because short-read data lacks INDELS at these sites, it is unlikely that any of these homopolymer deletions are real in these experiments. Dimer runs – stretches of repeating 2-mers (pronounced “two-mers”) – proved challenging regardless of basecaller. Mapping as above may be used to aid in manually calling these when they occur. Albacore is currently deprecated, and current versions of Guppy now incorporate a version of FlipFlop called Guppy High-Accuracy (HAC). Guppy HAC and subsequence versions were not evaluated in this work. Polishing is becoming standard practice for processing assemblies from ONT data because it redresses most homopolymer errors propagated into long-read-only assemblies. The best manually finished and polished contig had 1 error out of 16,722 bases, illustrating the utility of ONT hardware when paired with burgeoning software.

## Conclusions

HIV informatics, the study of HIV sequence information, has been limited by the common assumption that sequence fidelity exists between reference genomes available in sequence databases and similarly named HIV clones. Modern DNA sequencing methods, such as long- and short-read sequencing, are available to redress this issue. Long-read sequencing fills in gaps left behind by short-read interrogation of HIV-1. Current limitations of the approaches used in the present work to study HIV are 1.) the cost of long-read sequencing, regardless of platform, compared to the cheaper short reads from sequencing-by-synthesis, 2.) long DNA extraction methods in diseased tissue (Gener, unpublished), and 3.) the lower per-base accuracy (low-mid 90’s with ONT vs. 98-99% with ILMN or newer PacBio HiFi), including difficulty near homopolymers and dimer runs (**Supplemental Figure 5**). A nontrivial but redressable limitation is availability of personnel trained to prepare sequencing libraries, to run sequencing, and to analyze results. As the price of long-read sequencing decreases, hardware and software used in basecalling and library protocols improve, and with the advent of more user-friendly tools, the cost of obtaining usable data from long reads will become negligible compared to the ability to answer historically intractable questions. This work raises the possibility of being able to detect at least some recombination events, in a reference-free manner requiring only the comparison of LTRs from the same integrants (**Figure 6**). We suggest that pHXB2_D and pNL4-3 constructs may be used as negative and positive controls for the development of such screens. While other HIV reference proviral clones were reported to have identical LTR pairs, this remains to be tested in other clones, since other clones were generated with shorter sequencing methods. For example, pNL4-3_gag-pol(Δ1443-4553)_EGFP had distinct LTRs as a plasmid. However, if an NL4-3 virus is made from pNL4-3, the LTR sequences would homogenize to pNL4-3’s 3’ LTR sequence. Future work will include optimizing DNA extraction protocols with the goal of capturing higher-coverage fuller glimpses of each HIV proviral integration site in *in vivo* HIV models and patient samples. This work has broad implications for all cells infected by both integrating and non-integrating viruses, and for the characterization of targeted regions in the genome which may be recalcitrant to previous sequencing methods. Long-read sequencing is an important emerging tool defining the post-scaffold genomic era, allowing for the characterization of anatomical landmarks of hosts and pathogens at the genomic scale.

**Figure 6:**
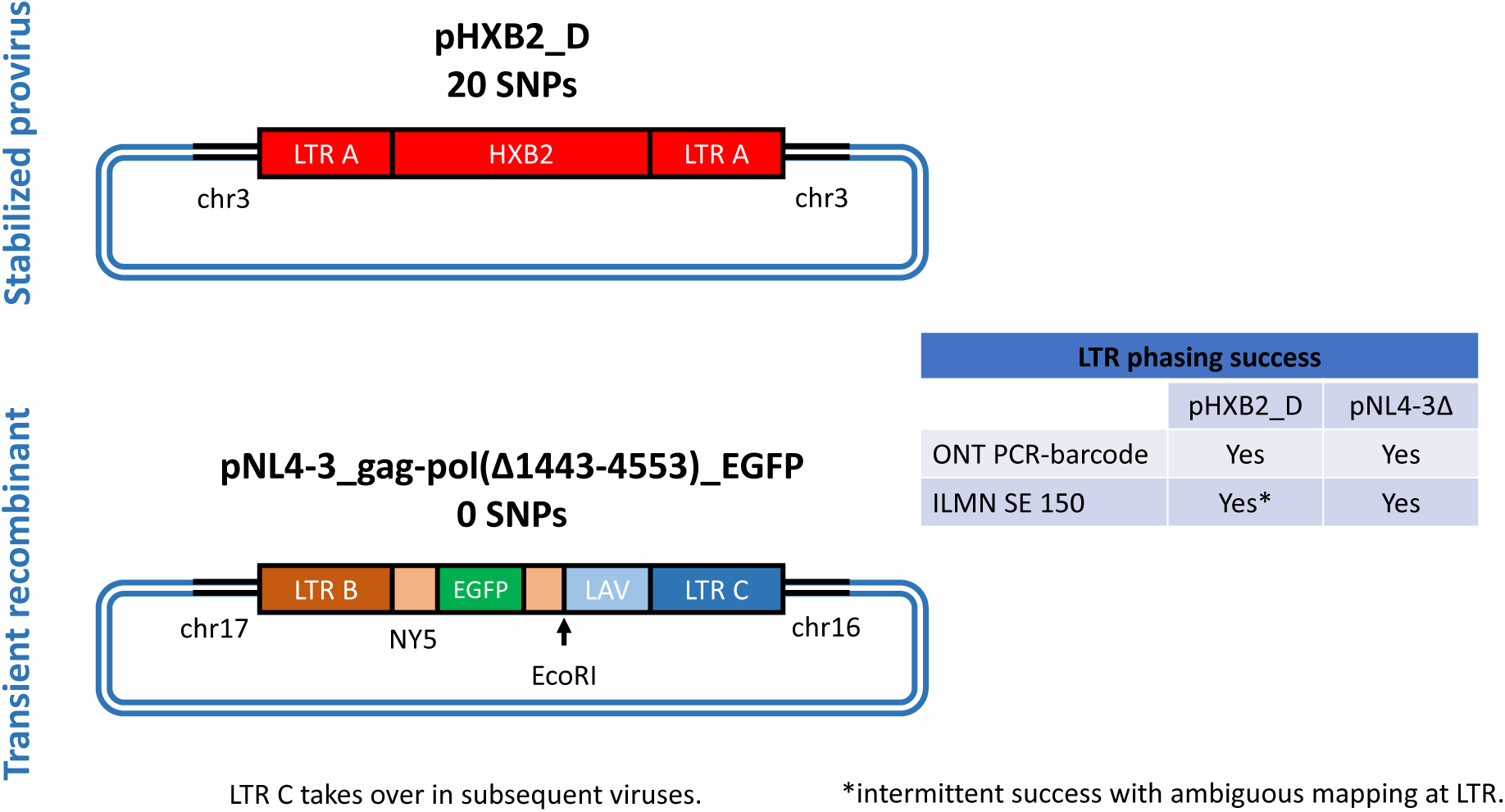
Summary of long-vs. short-read mapping by ability to phase LTRs.

## Disclaimer

Erratum: Preprint version 1 of this work [43] incorrectly cited the Integrated Genome Browser for work that was completed with the Integrative Genomics Viewer. Apologies for the mistake.

## Authors’ contributions

ARG conceived of this project, performed experiments, analyzed results, and drafted the manuscript. WZ rederived pNL4-3_gag-pol(Δ1443-4553)_EGFP. All authors discussed data and edited the manuscript. ARG and PK provided funding.

## Acknowledgements

As part of a summer bioinformatics internship in the Paul E. Klotman Laboratory at Baylor College of Medicine, Akash Naik supervised by ARG performed *in silico* mapping analyses/experiments, generated and/or aided in the synthesis of **Supplemental Figure 4**, and assisted in writing relevant portions, discussing, and editing this manuscript. During a second summer internship with American Physician Scientists Association Virtual Summer Research Program, the following students were supervised by ARG helped to create **Figure 1A** and **Supplemental Table 1**: Yini Liang, Kirk Niekamp, Maliha Jeba, Delmarie M. Rivera Rodríguez. Orthogonal sequence verification was performed as a service by staff at the Center for Computational & Integrative Biology DNA Core at Massachusetts General Hospital, Boston, MA, USA.

We would like to thank the staff at the DNA Core for their exceptional services, including expert analyses and rapid turnaround time. We would like to thank Drs. Steven Richards, Qingchang Meng and the staff of the Human Genome Sequencing Center Research (HGSC) and Development (R&D) team for their earlier support in nanopore adoption. We would like to thank the team at Oxford Nanopore Technologies for their timely improvements and continued R&D. I would also like to thank Ms. Taneasha Monique Washington (current) and former members of the Paul E. Klotman lab, Dr. Gokul C. Das and Alexander Batista. I would like to thank Dr. Alana Canupp and the late Dr. Jim Maruniak for their early interest in my scientific development, and for the passion that they show in everything that they do.

## Available additional files

Albacore basecalled barcode 10

Guppy basecalled barcode 10

FlipFlop basecalled barcode 10

Albacore basecalled barcode 11

Guppy basecalled barcode 11

FlipFlop basecalled barcode 11

Minimap2 and BWA-MEM alignments (.bam and .bai)

Clipboards from points of interest (verified SNVs; n=20)

.dna files of contigs (n=6)

MGH data (raw + contig)

Supplemental Tables

Supplemental Figures

## Supplemental Information

### Data exploration with long- and short-read mapping

To assemble pHXB2_D, we tried the following short read assemblers on short-read data from the external core: IDBA [46], MIRA [47], [48], SPAdes [49], and SSAKE [50], [51]. These were chosen as a convenience because they were already stably implemented in Galaxy (specifically usegalaxy.eu). Of these, SSAKE produced discontinuous assemblies with default parameters. The discontinuous contigs did however map to the core’s assembly (not shown).

### Enabling STEM outreach

This work was performed as two control experiments with identically prepared libraries for a STEM outreach initiative, Student Genomics (Gener, et al., manuscript in prep). Given the constraints of the Student Genomics pilot, a rapid sequencing kit with tagmentation (explained below) with PCR barcoding was used to pool samples for ONT sequencing, with the consequence of fragmenting plasmid DNA more than what would have been ideal for capturing full-length HIV. That said, these controls could have been just as easily replaced by any samples/experiments benefiting from long-read sequencing at moderate-to-high coverage.

### Supplemental Tables

**Supplemental Table 1: HXB2 is still a common HIV clone.**

See **Supplemental Digital Content.**

See also **Figure 1A.**

**Supplemental Table 2: HIV provirus clones**

See **Supplemental Digital Content.**

Of the HIV clones available through ARP, the table represents the only validated proviruses with both upstream and downstream homology arms mapping to the same integration sites. pNL4-3 is included as a known chimera with two integration half-sites. Other clones were made with cDNA cloning, usually TA cloning (per ARP entries). Note: Reference hg38. Aligner: minimap2 with “Long Assembly” mapping settings. All homology arms had Alignment quality = 60. Upstream = host plus strand; independent of integration orientation. Coordinates reported from UCSC. ARP = NIH AIDS Reagent and Reference Program. IS = integration site.

**Supplemental Table 3:** Variation in assemblies at the feature level. Assembled with Canu. NA denotes features which may not have matched exactly, but which were collapsed with adjacent features to facilitate counting. Variants called manually by mapping assemblies over HXB2 features with SnapGene.

|  | Mismatches |  |  |  |  |  | Gaps (INDEL) |  |  |  |  |  |
| --- | --- | --- | --- | --- | --- | --- | --- | --- | --- | --- | --- | --- |
|  | Taq |  |  | LA Taq |  |  | Taq |  |  | LA Taq |  |  |
|  | Albacore | Guppy | FlipFlop | Albacore | Guppy | FlipFlop | Albacore | Guppy | FlipFlop | Albacore | Guppy | FlipFlop |
| 5' LTR | NA | 9 | 9 | 9 | 9 | 9 | NA | NA | 0 | 2 | 0 | 0 |
| gag | 2 | 2 | 2 | 2 | 2 | 2 | 12 | 10 | 9 | 9 | 8 | 8 |
| 5' LTR+ψ | 10 | 10 | NA | NA | NA | NA | 5 | 2 | NA | NA | NA | NA |
| pol | 7 | 6 | 6 | 6 | 6 | 6 | 26 | 22 | 9 | 18 | 11 | 10 |
| vif | 0 | 0 | 0 | 0 | 0 | 0 | 3 | 3 | 2 | 3 | 1 | 1 |
| vpr | 0 | 0 | 0 | 0 | 0 | 0 | 1 | 1 | 1 | 0 | 0 | 0 |
| tat | 2 | 1 | 1 | 1 | 1 | 1 | 10 | 6 | 3 | 7 | 4 | 5 |
| rev | 2 | 1 | 1 | 1 | 1 | 1 | 10 | 7 | 4 | 7 | 4 | 5 |
| vpu | 1 | 0 | 0 | 0 | 0 | 0 | 0 | 0 | 0 | 0 | 0 | 0 |
| gp160 | 2 | 1 | 1 | 1 | 1 | 1 | 11 | 7 | 4 | 8 | 4 | 5 |
| nef | 1 | 1 | 1 | 1 | 1 | 1 | 3 | 2 | 2 | 3 | 2 | 1 |
| 3' LTR | 2 | 2 | NA | NA | NA | NA | 2 | 0 | NA | NA | NA | NA |
| nef+3' LTR | NA | 2 | 2 | 2 | 2 | 2 | NA | 2 | 2 | 5 | 2 | 1 |
| HXB2 | 22 | 20 | 20 | 20 | 20 | 20 | 61 | 46 | 28 | 47 | 27 | 25 |
| Downstream bridge | 0 | 0 | 0 | 0 | 0 | 0 | 4 | 5 | 1 | 2 | 1 | 1 |
| pBR322-related | 0 | 0 | 0 | 0 | 0 | 0 | 19 | 19 | 13 | 18 | 19 | 15 |
| Upstream bridge | 2 | 2 | 3 | 2 | 2 | 2 | 8 | 6 | 5 | 7 | 7 | 3 |

### Supplemental Figures

**Supplemental Figure 1A:**
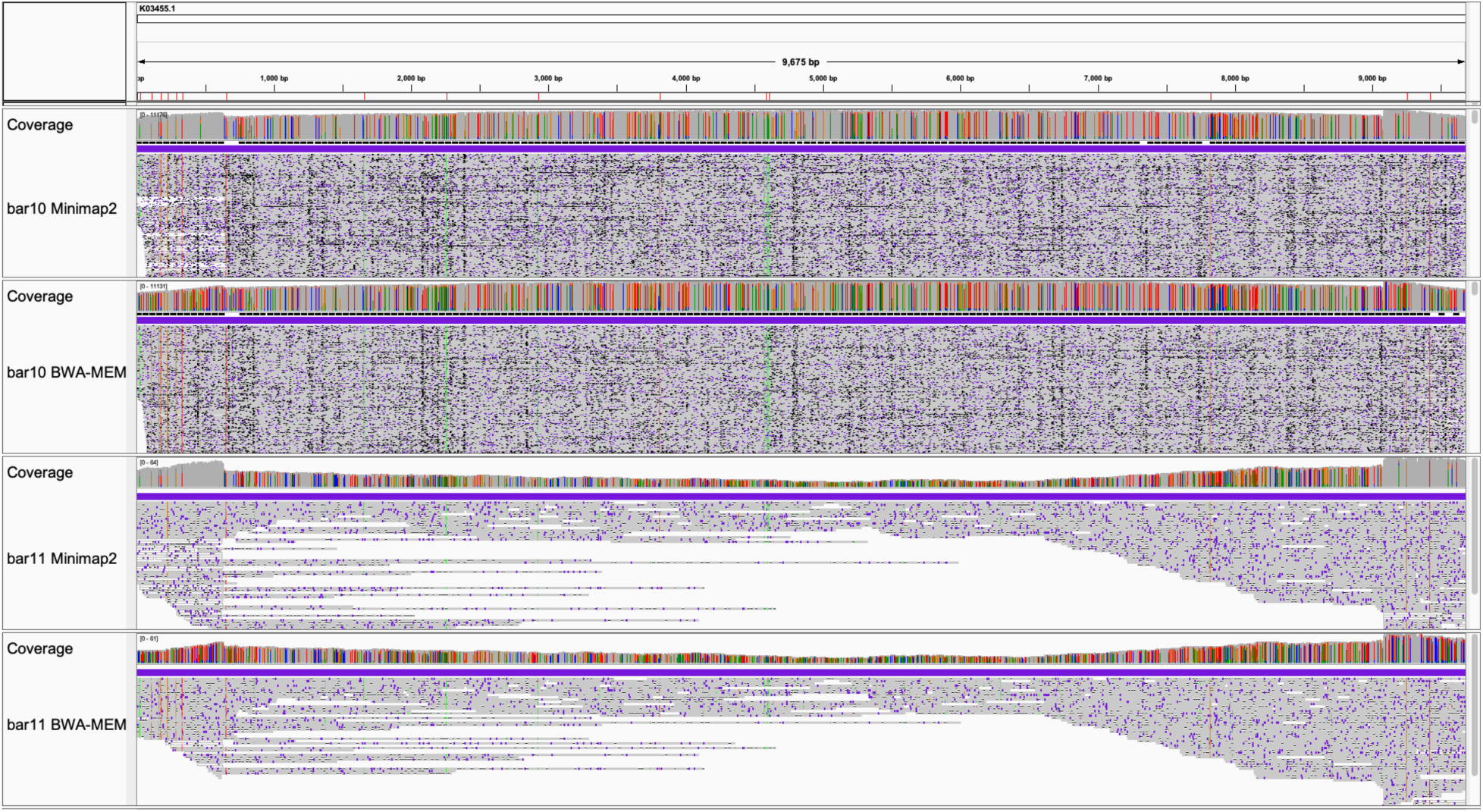
Unbiased nanopore DNA sequencing coverage over HXB2 depends on DNA polymerase and mapper. ONT basecaller = Albacore (worst).

**Supplemental Figure 1B:**
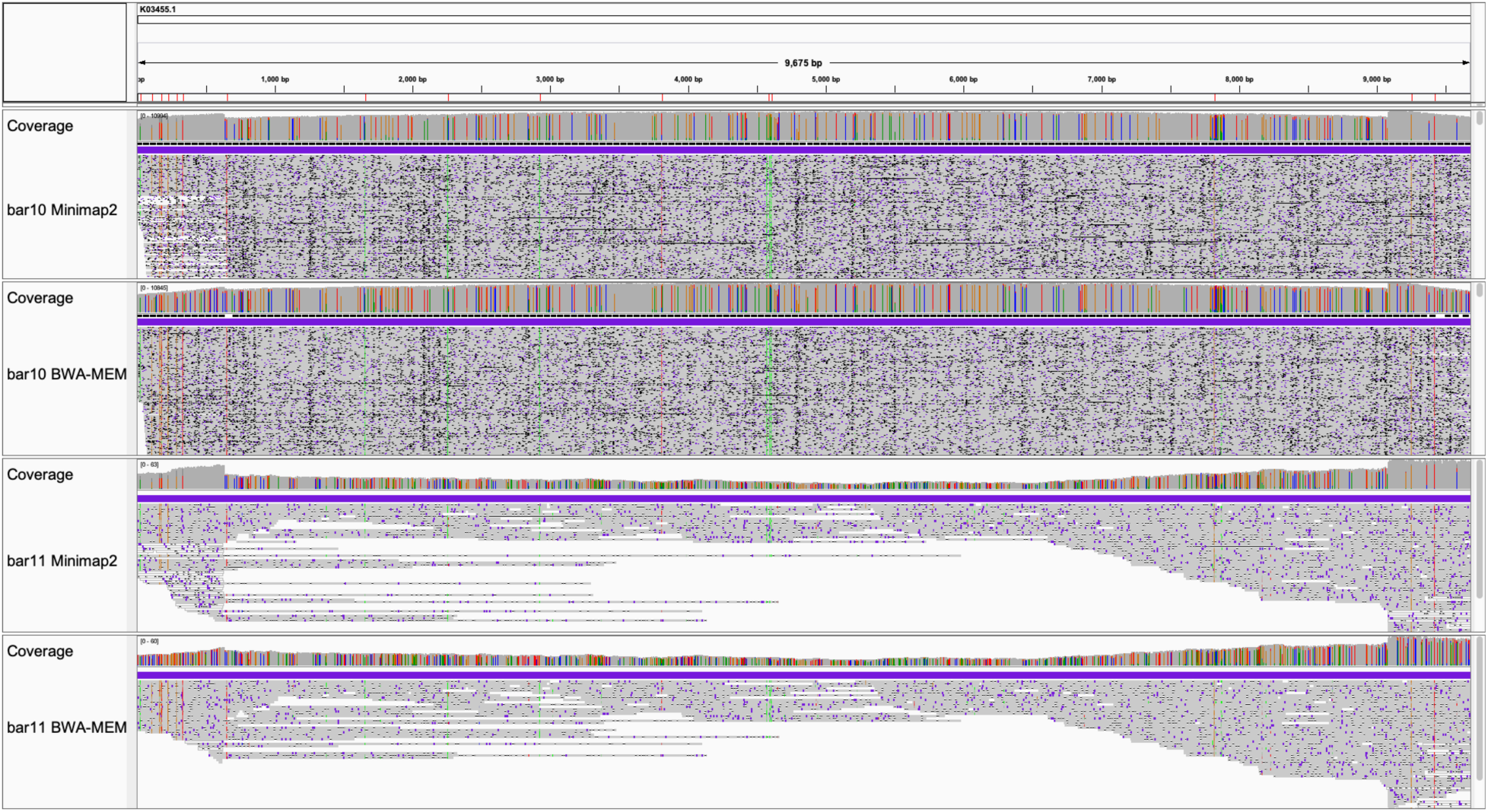
Unbiased nanopore DNA sequencing coverage over HXB2 depends on DNA polymerase and mapper. ONT basecaller = Guppy.

**Supplemental Figure 1C:**
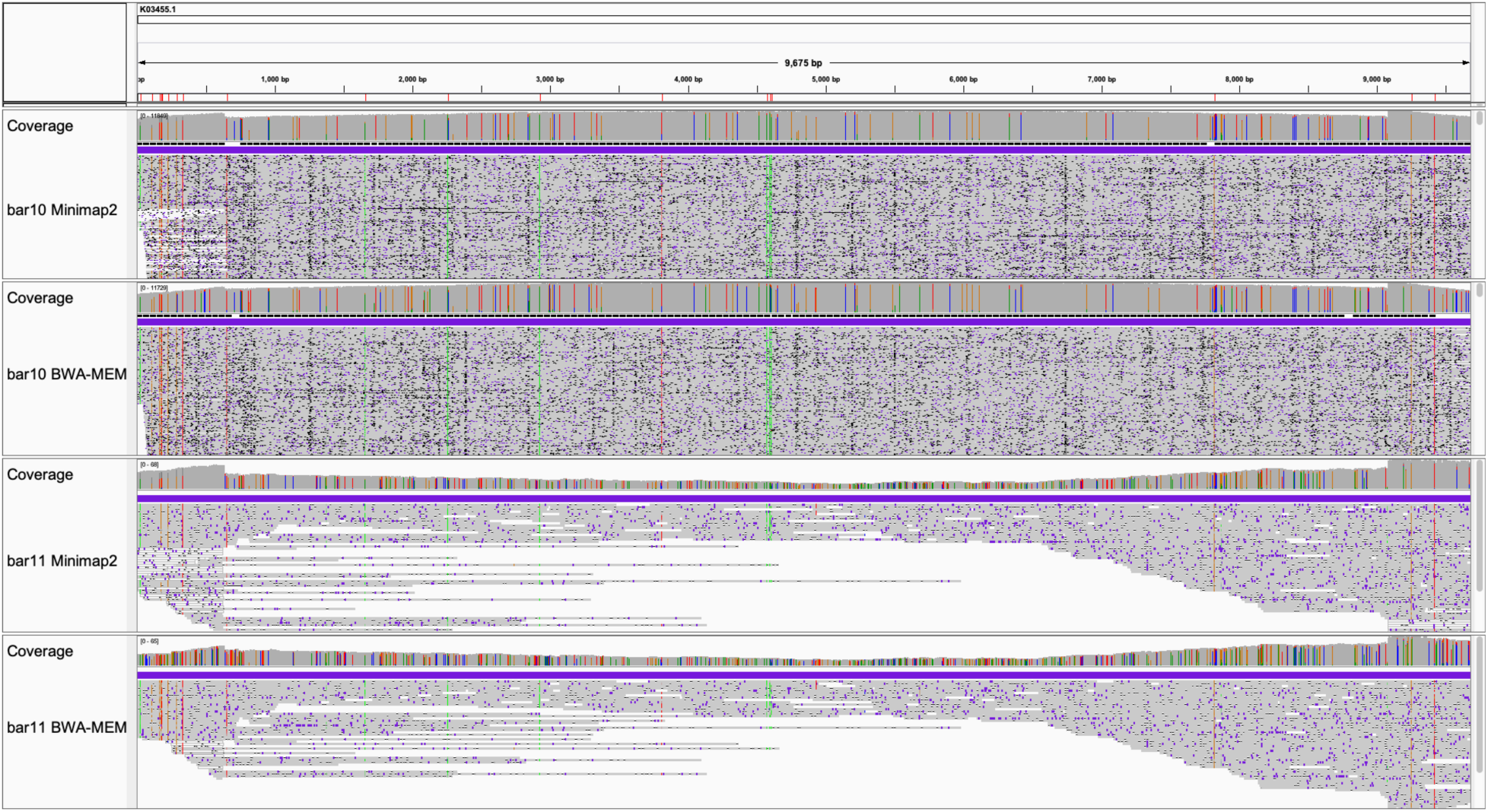
Unbiased nanopore DNA sequencing coverage over HXB2 depends on DNA polymerase and mapper. ONT basecaller = FlipFlop (best). Top two Coverage and Alignment panels from barcoded library 10 (bar10 = LA Taq). Bottom two from Barcode 11 (bar11 = Taq). Minimap2 and BWA-MEM were used to map reads basecalled with Albacore (worst), Guppy, or FlipFlop (best) to HXB2. Color-coding: Red below genome scale marks 20 SNVs across the HIV segment of pHXB2_D. Purple is an insertion in a given read relative to reference. White is either a deletion in a given read or space between two aligned reads. Gray in alignment field means base same as reference, and in coverage field means major allele is at least 95% the same as reference. Per-read “insertions” and “deletions” do not necessarily represent true insertions or deletions actually present in the sample, because each read is likely an imperfect independent observation. Automated assembly followed by manual consensus building converts these overlapping reads into approximations of the ground truth. “Unbiased” refers to not amplifying a given region (e.g., pol) before ligating ONT sequencing adapters. In the present approach, the tagmentation process randomly cuts DNA, creating ∼2000 bp pieces. Tagmented DNA is then amplified based on tagmentation adapters.

**Supplemental Figure 2:**
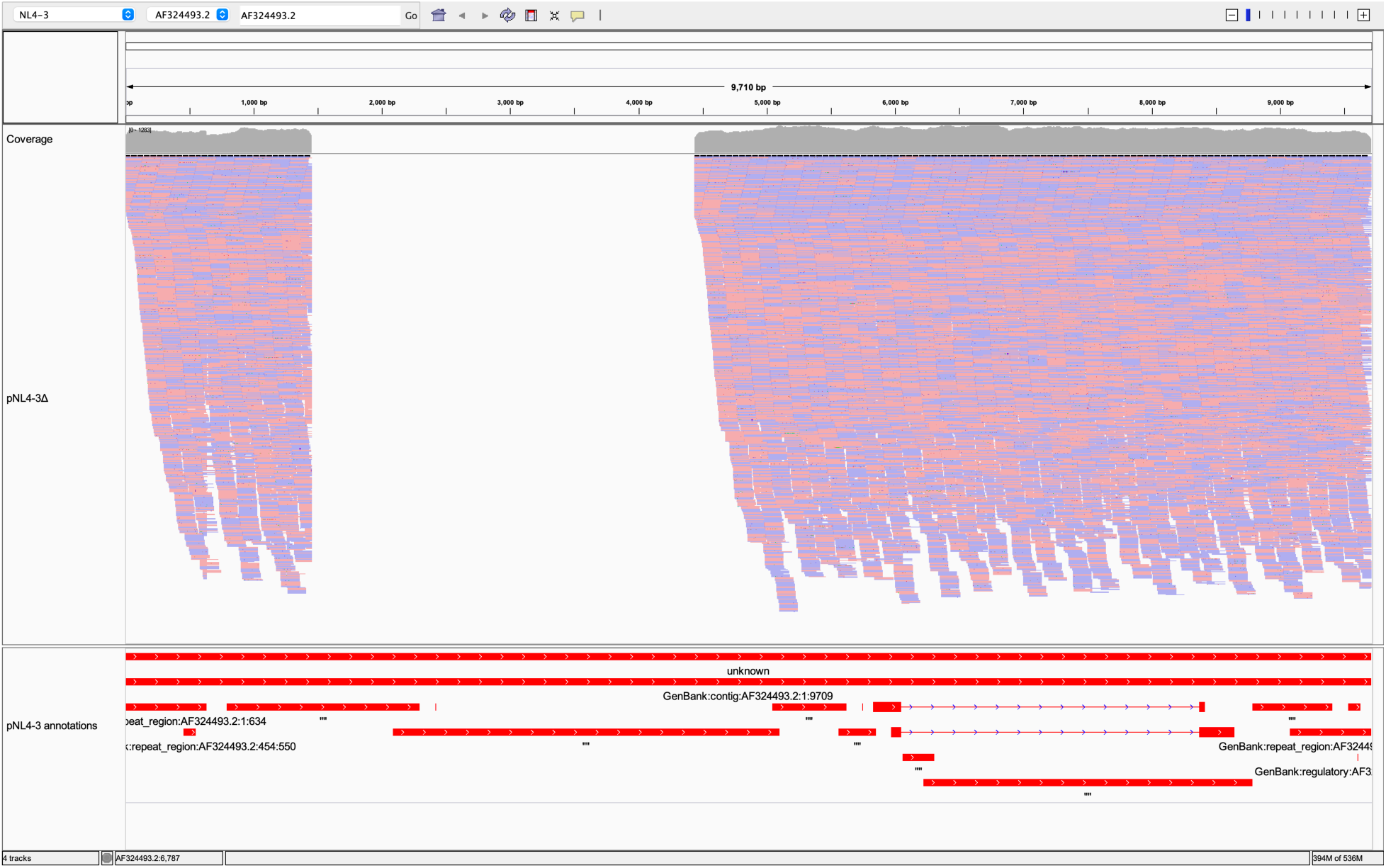
Reads map well to HIV-1 NL4-3 segment of pNL4-3 assembly because NL4-3 LTRs are distinct.

**Supplemental Figure 3A:**
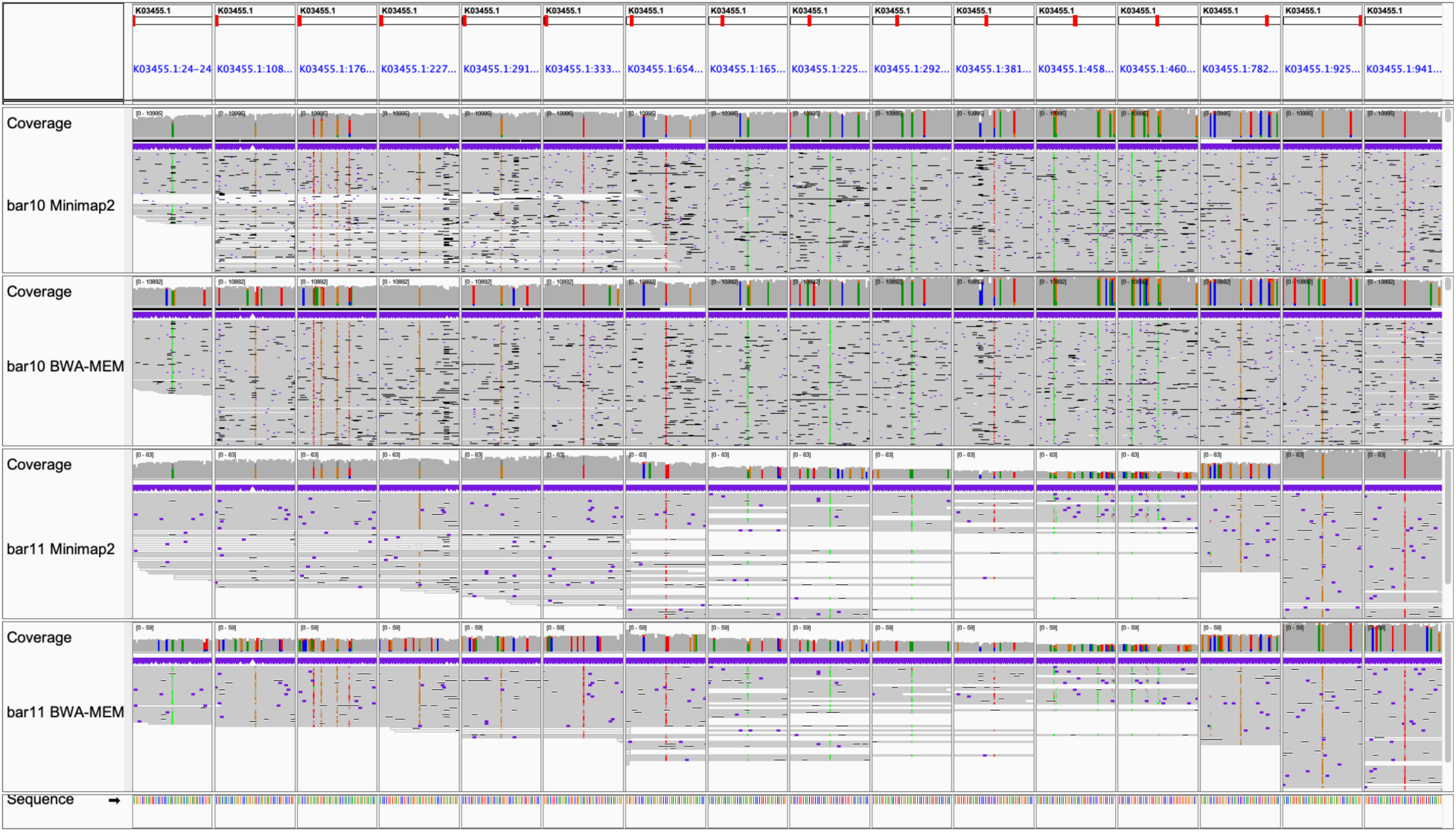
HIV single nucleotide variants (SNVs) in pHXB2_D. ONT basecaller = Albacore (worst). Gray indicates per-base consensus accuracy ≥ 80%. These alignments are the noisiest (less gray and most divergent from reference) between Supplemental Figures 3A, 3B, and 3C.

**Supplemental Figure 3B:**
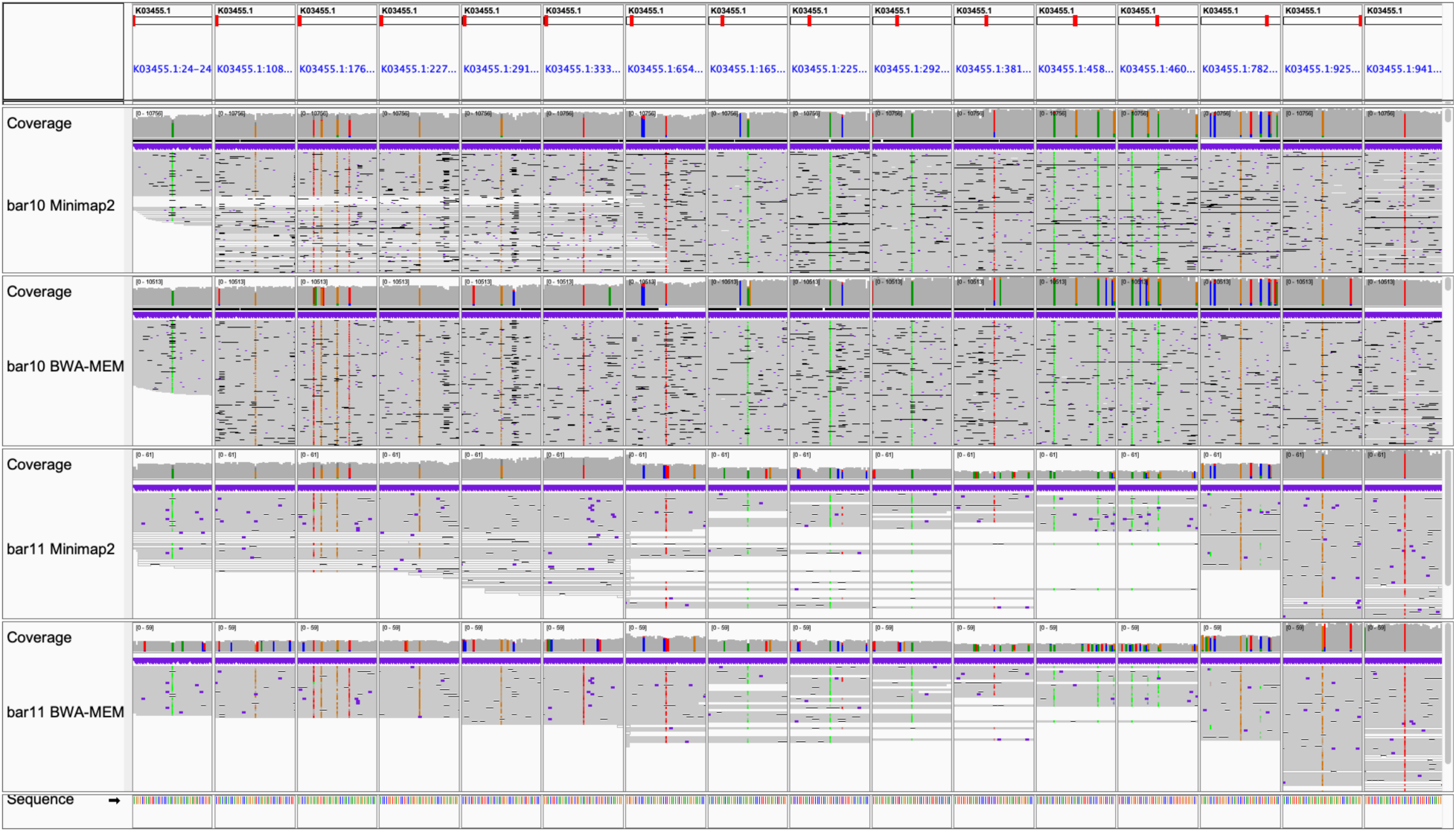
HIV single nucleotide variants (SNVs) in pHXB2_D. ONT basecaller = Guppy.

**Supplemental Figure 3C:**
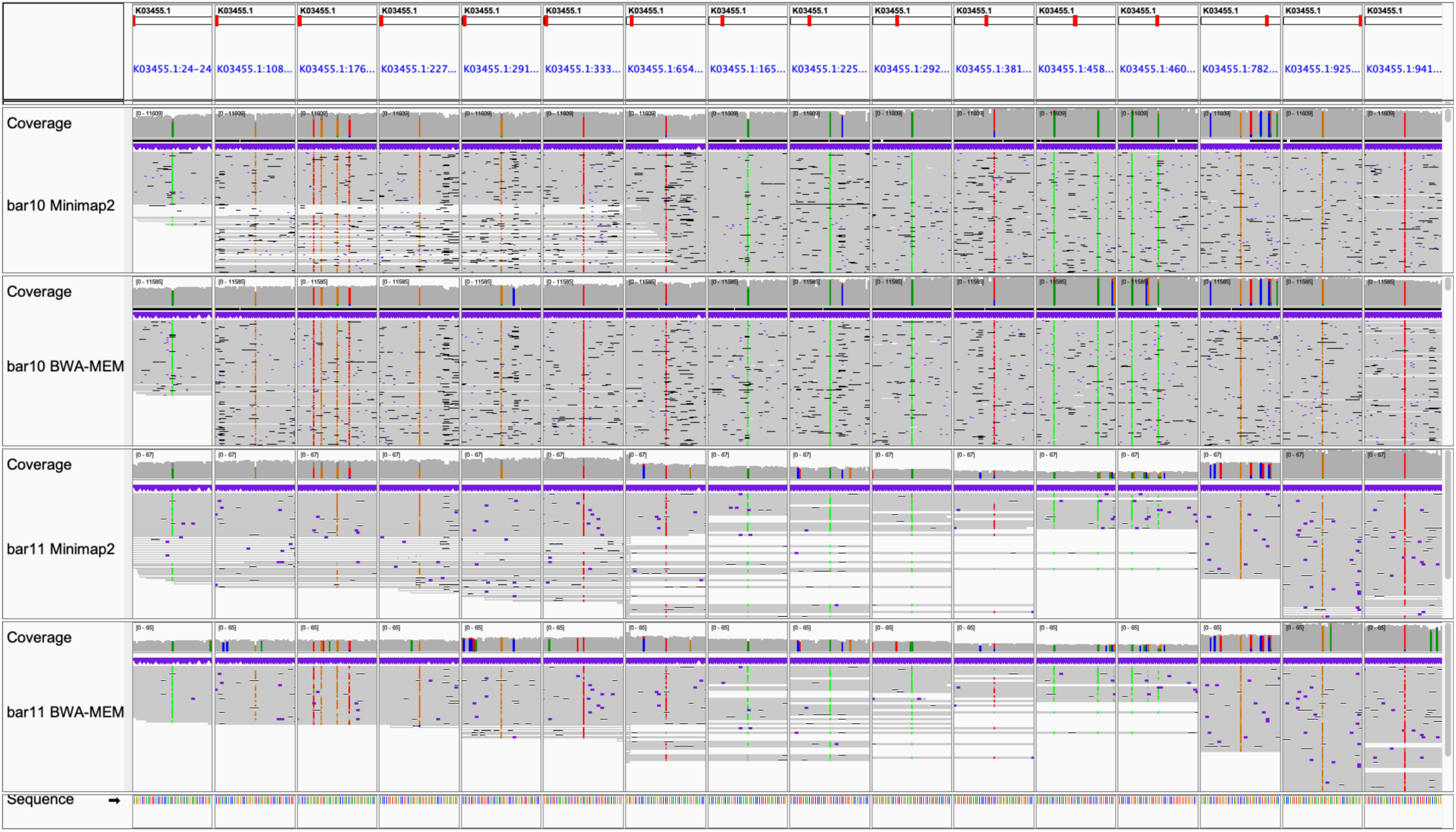
HIV single nucleotide variants (SNVs) in pHXB2_D. ONT basecaller = FlipFlop (best). These alignments are the least noisy (most gray and like reference) between Supplemental Figures 3A, 3B, and 3C.

**Supplemental Figure 3D:**
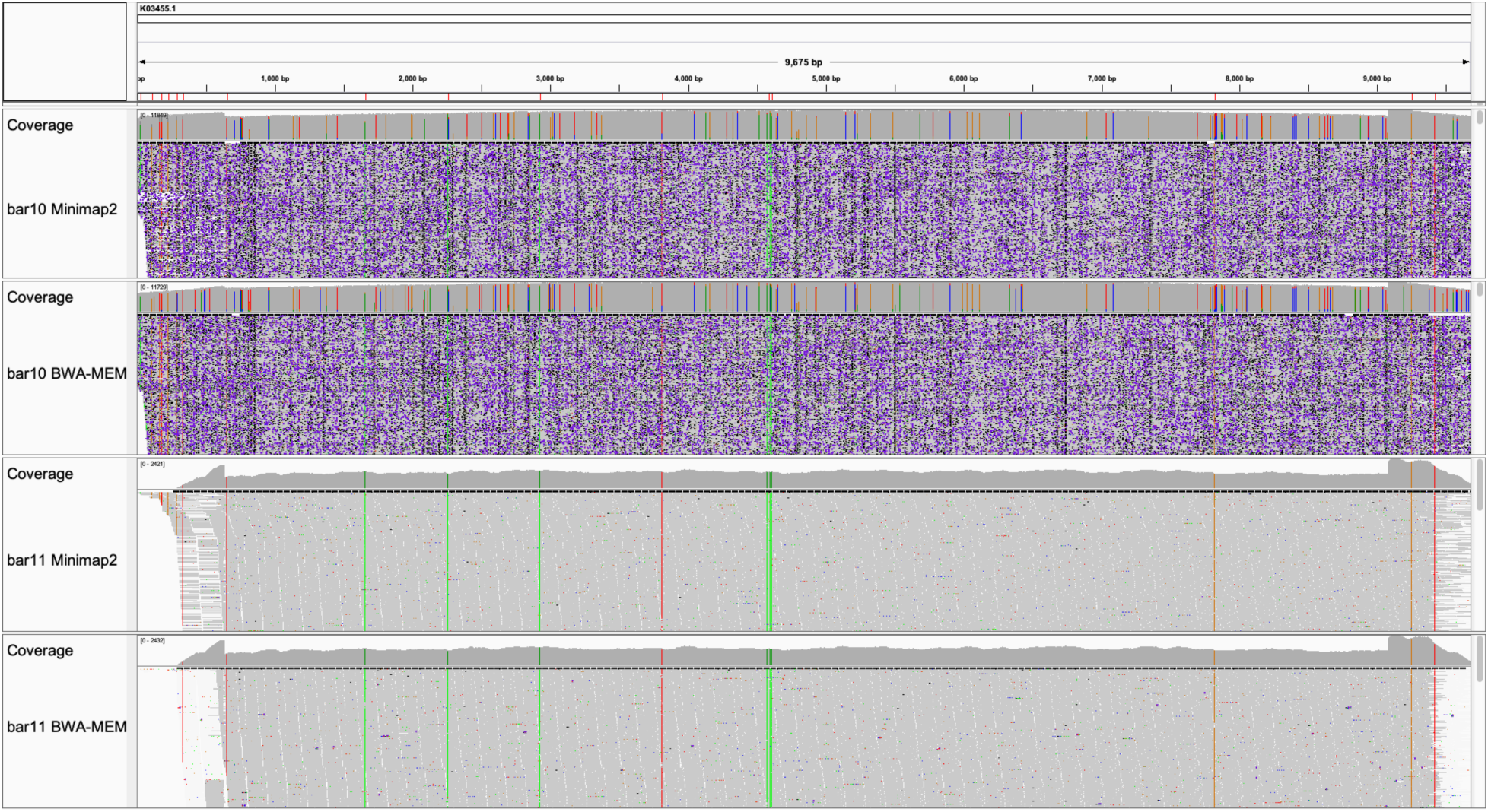
HIV single nucleotide variants (SNVs) in pHXB2_D, long vs. short reads (HIV genome). Long reads outperform short reads at HIV-1 LTRs. ONT basecaller = FlipFlop. Short read as single-end 150, clipped to 142, provided by external core. Mappers = Minimap2 (better), BWA-MEM.

**Supplemental Figure 3E:**
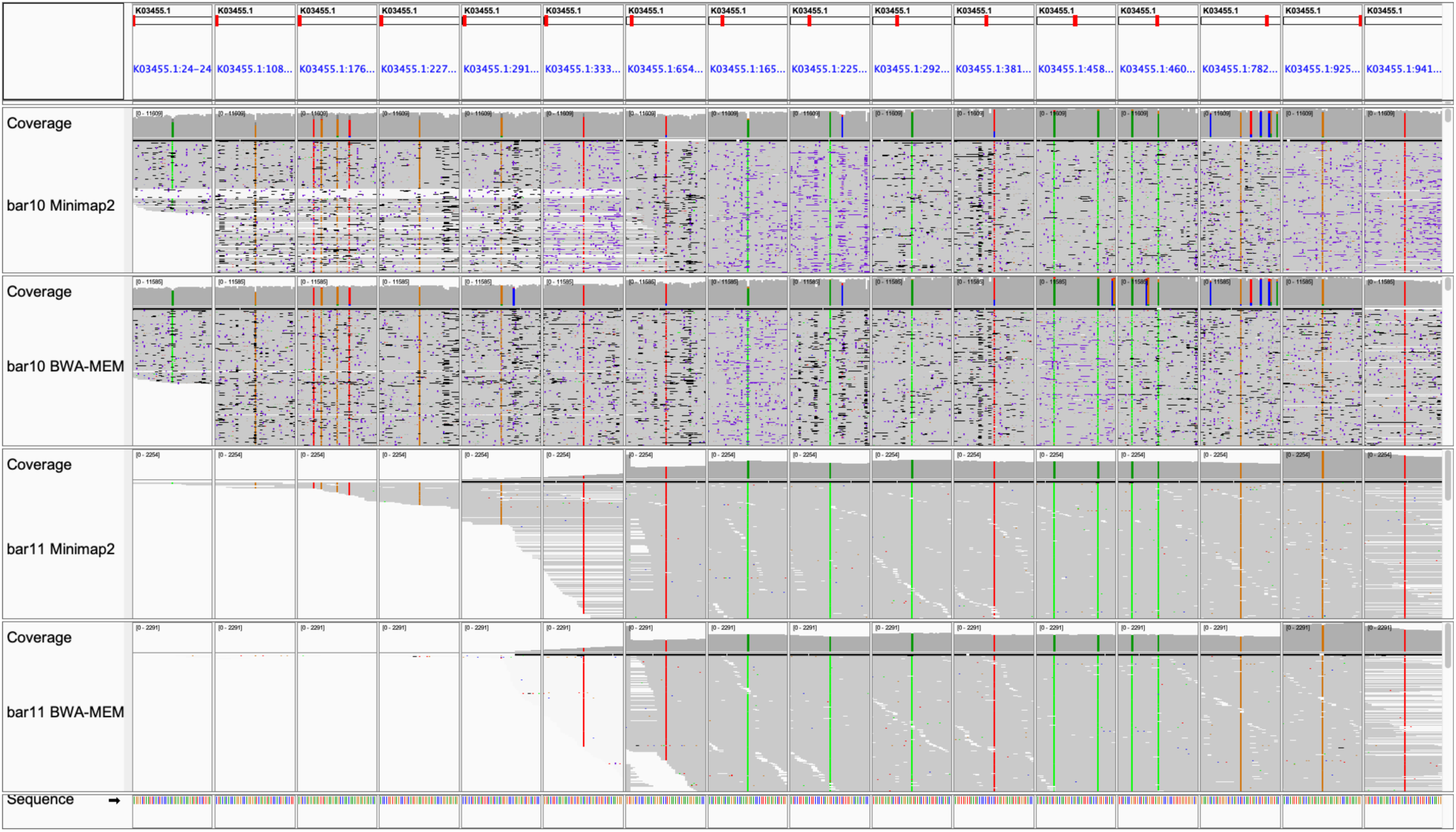
HIV single nucleotide variants (SNVs) in pHXB2_D, long vs. short reads (20 SNV-focused).

**Supplemental Figure 4A:**
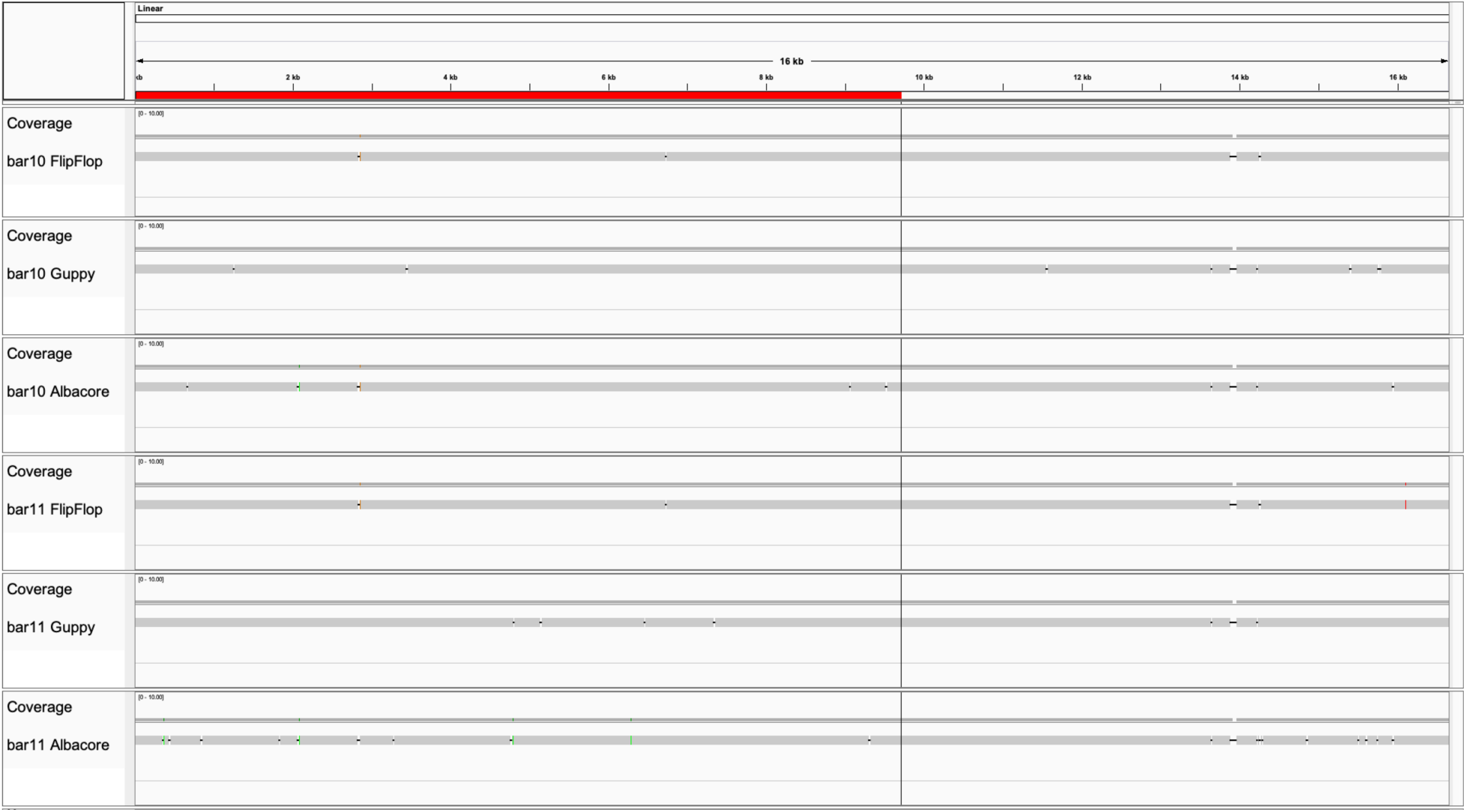
Assembling pHXB2_D from long reads only, varying basecaller and polymerase. Each pane (n=6) summarizes the results of contig curation. Divergence from reference decreases with newer basecallers, and with long amplicon DNA polymerase (Sigma-Aldrich Taq vs. LA Taq by Takara). Errors in assembly occurred at homopolymers (most often deletions not visible at this resolution; see **Supplemental Figure 6**), dimer or trimer runs. bar10 = LA Taq library. bar11 = Taq library. pHXB2_D Genbank:MW079479. Best contigs presented, manually curated to match pHXB2_D coordinates. Note LTRs (beginning and terminal 634 bp of red bar) are resolved in almost all assemblies. See **Supplemental Table 3** for differences between assemblies and the reference (left red). Plasmid backbone (right) differences are not reported.

**Supplemental Figure 4B:**
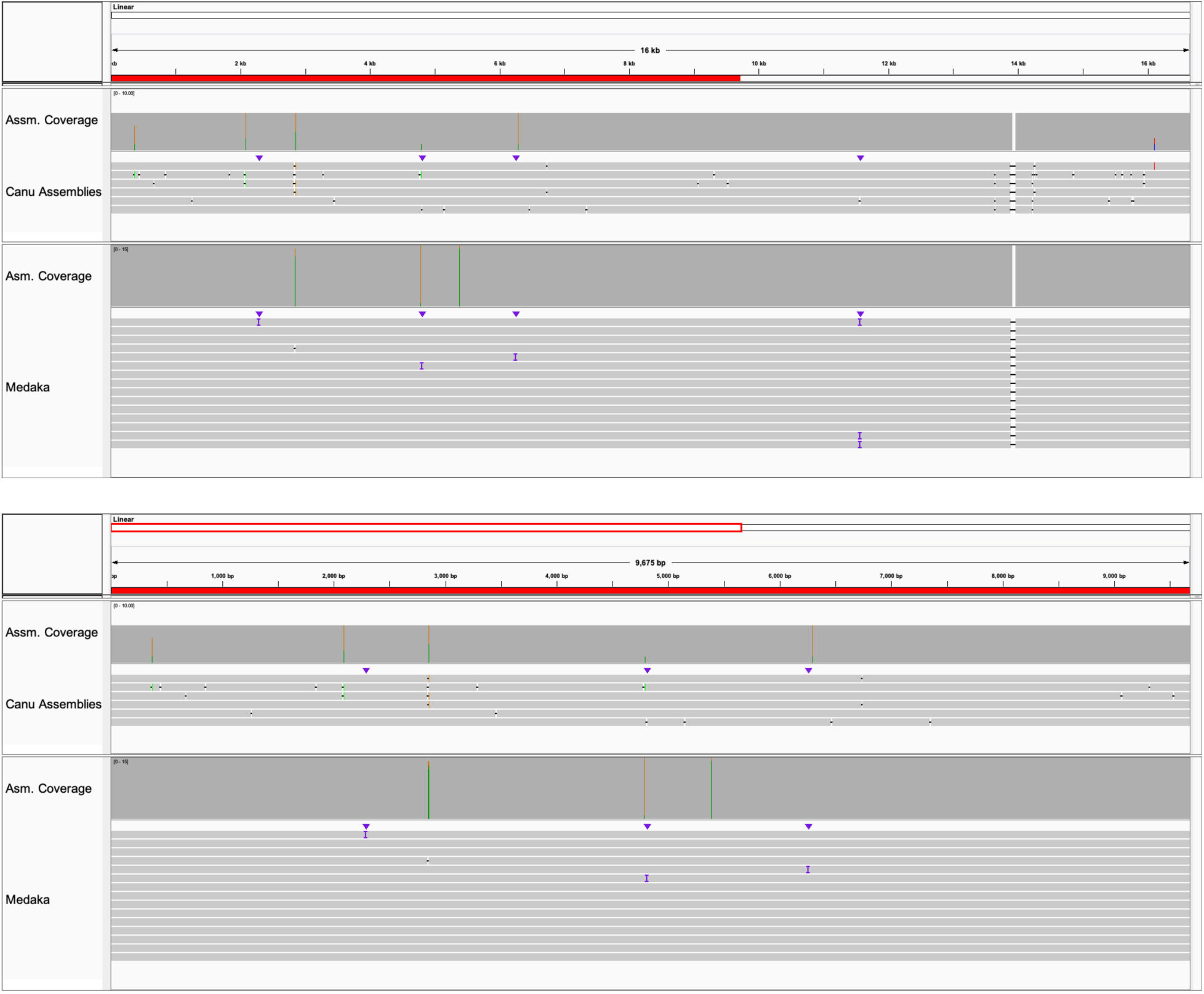
ONT errors corrected by polishing ONT-only assemblies. Assemblies polished with Medaka (ONT). Top: pHXB2_D genome. Bottom: HIV-only segment. The best polished assembly had one error in the entire plasmid (1 error out of 16,722 bases), with a corresponding consensus accuracy of 99.99402%. This happened to be in HIV segment (HIV-1 between position 1 and 9719; 1 error out of 9719 bases), with corresponding accuracy of 99.989711%. Note the conserved 52 bp gap in the backbone of pHXB2_D was redundant sequence included in the short-read assembly from the core. It was not supported by long-read data, and therefor was validated as a technical artifact from the core’s pipeline. Reference: short-read assembly. LTRs (beginning and terminal 634 bp of red bar) are resolved in polished assemblies.

**Supplemental Figure 4C:**
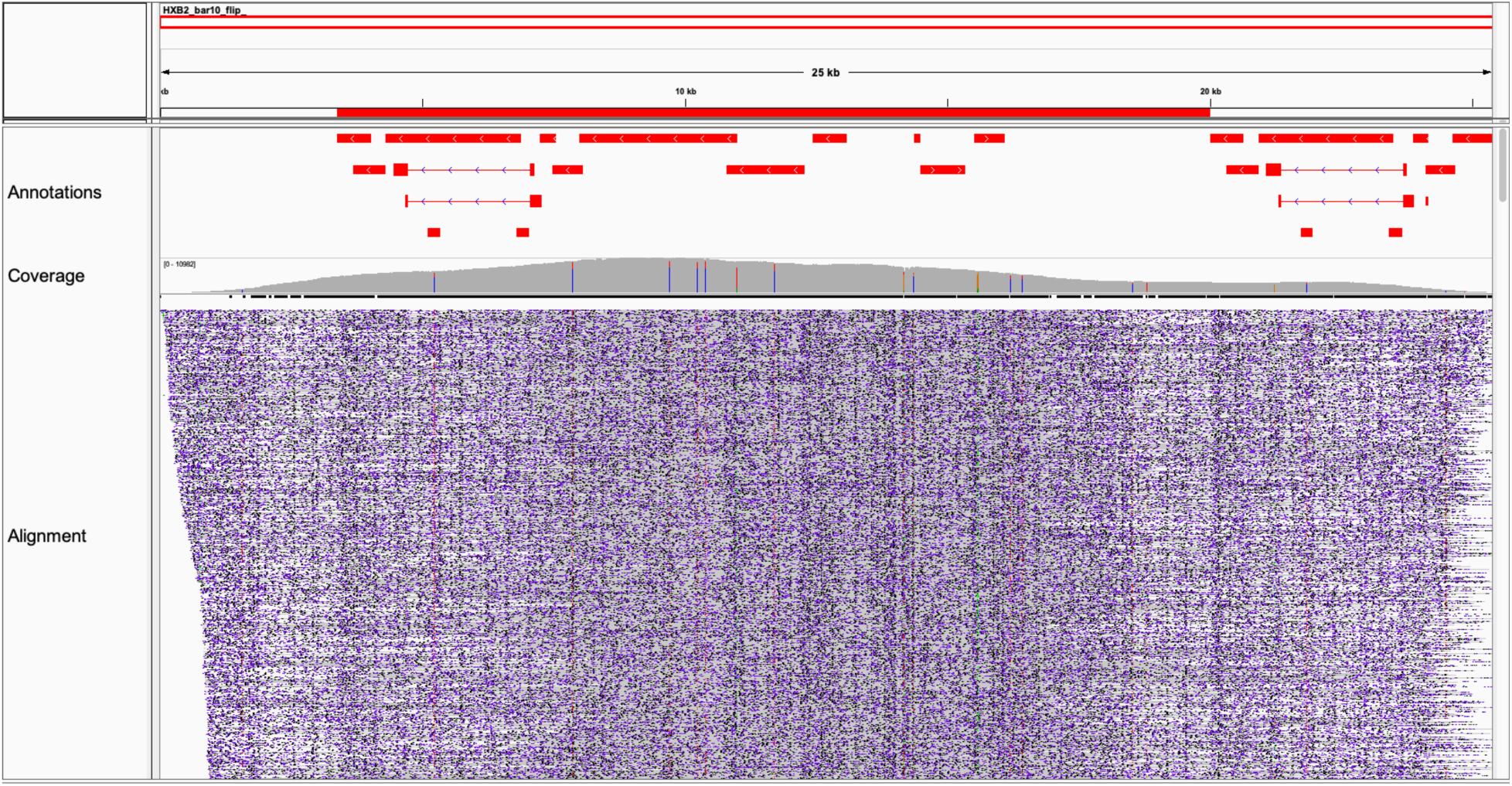
Mappability of long reads over contigs during assembly quality control. Coverage depends on context. Abrupt changes in coverage from terminal regions of HXB2 (**Figure 1F, Supplemental Figures 1 and 3**) were artifacts from supplying mappers with an HIV reference without a plasmid backbone. Long reads from barcode 10 (LA Taq) mapped with minimap2 [23] and “reference” contig from assembly (Canu v1.8) with basecalled data (FlipFlop). Stripes in this figure are not SNVs. They represent technical variability at homopolymers. Assemblies were manually curated to start with 5’ LTR in the sense orientation, leaving the plasmid backbone on the left. Because there were not real insertions in the HIV segment, the HIV coordinates are the same as HXB2 (both 9719 bp long). Compare with 52 bp technical artifact from the core’s short read assembly in **Supplemental Figure 4A, 4B**.

**Supplemental Figure 5:**
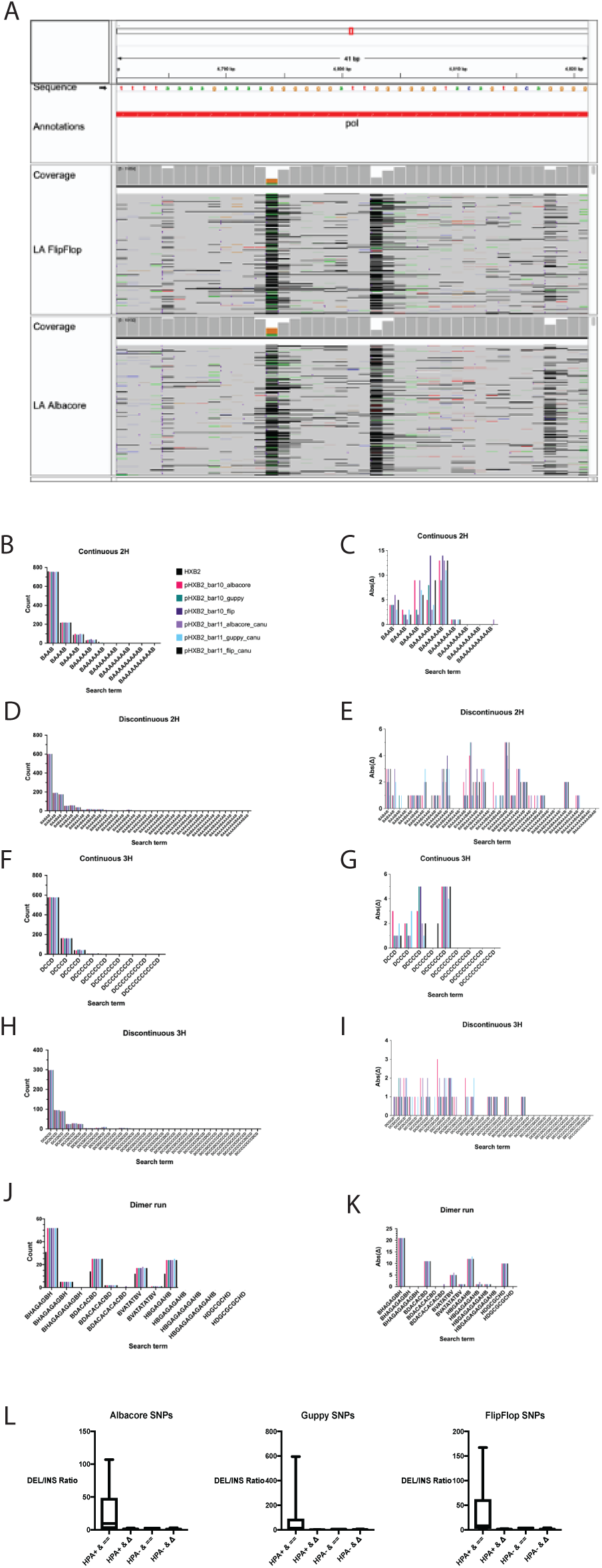
Homopolymers and dimer runs are ONT artifacts in unpolished assemblies. Supplemental Figure 5A: A set of homopolymer tracks from HXB2 plasmid. Alignments with BWA-MEM shown from FlipFlop (top) and Albacore (bottom) basecalled reads. Mapping is pre-assembly. Supplemental Figure 5B: Continuous 2H counts in unpolished assemblies. 2H = A or T homodimers. Supplemental Figure 5C: Continuous 2H Absolute Difference. Supplemental Figure 5D: Discontinuous 2H counts in unpolished assemblies. Supplemental Figure 5E: Discontinuous 2H Absolute Difference. Supplemental Figure 5F: Continuous 3H counts in unpolished assemblies. 3H = C or G homodimers. Supplemental Figure 5G: Continuous 3H Absolute Difference. Supplemental Figure 5H: Discontinuous 3H counts in unpolished assemblies. Supplemental Figure 5I: Discontinuous 3H Absolute Difference. Supplemental Figure 5J: Dimer run counts in unpolished assemblies. Supplemental Figure 5K: Dimer run Absolute Difference. Dimer runs as pairs are the most problematic, with runs as triplets being resolvable by ONT. Supplemental Figure 5L: The ratio of deletions to insertions is higher at mismatches both adjacent to homopolymers and similar to neighbor bases. Box plot shows median (“x” is mean) and quartile ranges. Y-axis is ratio. HPA: homopolymer-adjacent. ==: same as neighbor base. Δ: different than neighbor base. Higher coverage (above ∼10) usually makes up for current error profile. Above true for Albacore, Guppy, and FlipFlop.

**Supplemental Figure 6:**
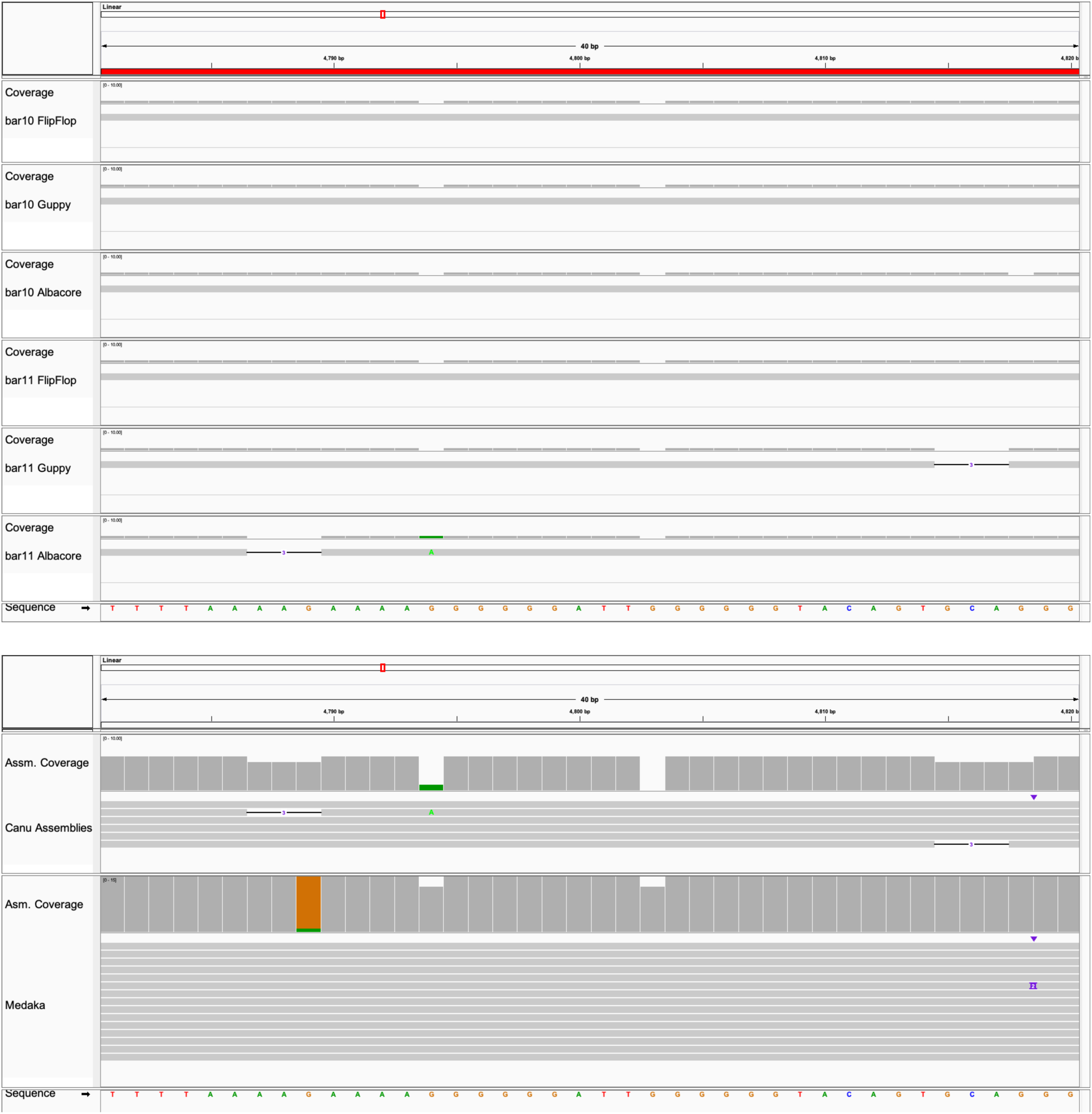
Assembly partially resolved homopolymers, which are improved by polishing. Top: Six ONT-only assemblies. Bottom: polished ONT-only assemblies, varying Medaka models. Deletions at 5’ of G homopolymers were not corrected, regardless of basecaller or Taq isoform. Note that polishing was not performed. IGV window is Linear:4,781-4,820. Bottom: polishing canu assemblies with medaka abrogated most ONT artifacts. Best medaka setting tested: r941_min_high_g330.

